# Effects of Electrocardiograms QRS Detection Algorithms in Heart Rate Variability Metrics

**DOI:** 10.1101/2025.04.23.650299

**Authors:** Alejandro Weinstein, Julio Rodiño, Mónica Otero

## Abstract

Heart Rate Variability (HRV) is a marker used for assessing autonomic nervous system function, derived from the timing between R-peaks in electrocardiogram (ECG) signals. Accurate detection of QRS complexes is essential for reliable HRV computation. While many R-wave detection algorithms exist, their impact on the accuracy of HRV metrics remains underexplored. This study addresses this gap by assessing how QRS detection errors affect HRV analysis across different algorithms and recording setups. We evaluated eight widely used QRS detectors using ECG recordings from 25 healthy participants under rest, cognitive load, and physical activity conditions. Two acquisition setups were considered: “chest strap” and “loose cables.” We used the manually annotated R-peaks to calculate the ground-truth HRV metric values. The relationship between the detector performance and HRV errors was evaluated for 11 metrics using the concordance correlation coefficient (CCC). Results showed significant variability in detector performance across algorithms and setups. No single QRS detection algorithm outperformed across all scenarios. Loose cable recordings yielded higher CCC values than chest straps, particularly for MeanNN and LF power. These findings highlight the critical role of QRS detector selection and signal acquisition conditions in HRV analysis. They underscore the need for context-specific benchmarking, particularly for wearable and ambulatory applications where signal quality can vary. Ultimately, this study offers practical recommendations for clinicians and researchers on selecting QRS detection algorithms that best align with their specific analytical objectives and recording conditions.

## I. Introduction

HEart rate variability (HRV) measures the variation in time intervals between successive heartbeats and reflects the autonomic nervous system’s (ANS) influence on the heart’s rhythm [1]. It is commonly used as an indicator of neurocardiac function, arising from heart-brain interactions and dynamic non-linear processes within the ANS [2]. HRV emerges due to interdependent regulatory mechanisms operating across various time scales, enhancing our ability to adapt to environmental stimuli and psychological stressors [3].

HRV is quantified using a wide range of time-domain, frequency-domain, and nonlinear metrics [4]. These measures serve as clinically relevant biomarkers for various diseases [5] and have been extensively used in psychophysiological research. For instance, time-domain metrics such as SDNN (standard deviation of NN intervals) and RMSSD (root mean square of successive differences) are commonly used to assess cardiovascular risk and monitor both acute and chronic heart conditions [4], [6], [7]. Frequency-domain features, including LF (low-frequency) and HF (high-frequency) power, have been found to differentiate patients with Type 2 Diabetes Mellitus [8]. Additionally, the LF/HF ratio, representing the balance between sympathetic and parasympathetic activity, has been studied in relation to depressive symptoms. For instance, research has shown that patients with depression and panic disorder exhibit a decreased LF/HF ratio, suggesting altered autonomic function associated with these conditions [9]. These diverse metrics provide insight into autonomic regulation and support their application across medical and psychological domains.

HRV metrics can be derived from various sources, including the electrocardiogram (ECG) [2], photoplethysmography [10], and facial videos [11]. Among these signals, the ECG is considered the gold standard for obtaining HRV metrics, as the R-peak component clearly signals the heartbeat [12]. Estimating RR intervals—the time between one R-peak and the next one—from ECG signals is crucial for accurately assessing HRV, as detailed in the following paragraph.

The process for obtaining a given HRV metric from the ECG can be split into three independent stages (see Fig. 1). The first stage is the data acquisition stage, where the raw ECG signal is recorded from the subject. This stage may include some signal preprocessing [13]. The output of this stage is the ECG signal. In the second stage, the ECG is the input to the QRS complex detector algorithm. The output of this stage is a vector [*T*_0_, *T*_1_, …], known as the interval tachogram or RR tachogram [3], [14], where its entries correspond to time differences between consecutive R-peaks.^1^ Note that although the objective of the QRS detector is to detect the QRS complex, in practice, these detectors use the R-peak as a fiducial point of the complex [13]. The input to the third stage is the interval tachogram, from which a given HRV metric is computed. As we can see, RR interval and HRV-based metrics computation are independent stages. Therefore, to compute a specific HRV metric, one must decide which QRS detection algorithm to use among all the existing ones (see Fig. 1).

**Fig. 1:**
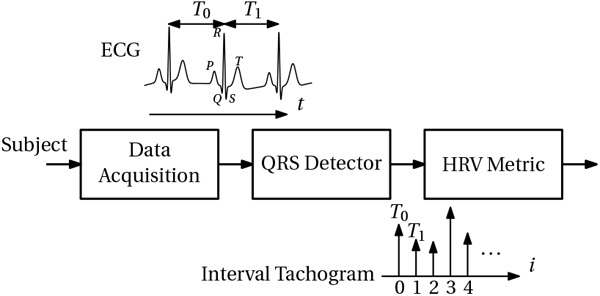
HVR metric computation pipeline: The data acquisition process records the raw ECG from a subject. After detecting the QRS complexes in the ECG, the time intervals *T*_0_, *T*_1_, … between consecutive R-peaks are determined. These sequences of intervals are used to build the interval tachogram. A given HRV metric is then computed from the interval tachogram.

HRV is a rate of a rate, thus, computing it involves a second derivative [16]. Since noise and high-frequency variations are amplified each time a derivative is computed [17], HRV is very sensitive to errors in the interval tachogram [18]. At the same time, different QRS detection algorithms have different performances (in terms of their precision, accuracy, etc.). The relationship between the method and its performance depends on the signal-to-noise ratio of the ECG [16], [19]. Moreover, the sensitivity to these errors depends on the specific HRV metric being computed [18]. For these reasons, this work studies how the selection of the QRS detection algorithm affects the errors in HRV metrics and how this relationship depends on the signal-to-noise ratio and artifacts in the ECG signal.

In previous works, Porr and Macfarlane [16] highlight performance differences among popular QRS detectors. The authors present a performance measure that combines temporal jitter with the F-score, called JF, to compare QRS detection algorithms. Their results show significant performance differences among the detectors as a function of the recording conditions. However, this work does not study how these differences may affect HRV metrics. The work of Lu et al. [20] is a systematic review of the different sources of uncertainties in HRV analysis. While it establishes that the ECG recording conditions and the QRS detection algorithm are significant sources of uncertainty in obtaining HRV metrics, it does not study how specific QRS detectors affect the errors in these metrics. Petelczyc et al. [18] studied the effects of errors in the interval tachogram on HRV metrics. They proposed a stochastic model for the interval tachogram. They used an empirical interval tachogram obtained using a single QRS detector, followed by a Monte Carlo simulation, to evaluate the effect of different noise levels. They did not consider different QRS detectors. Rohr et al. [21] performed a similar study using a more extensive stochastic model that considered not only Gaussian noise but also uniform, triangular, and t-student distributed noise. As before, they did not consider different QRS detectors.

To the best of our knowledge, our work is the first to investigate how different QRS detector algorithms affect the errors in HRV metrics. We used real ECG signals recorded under various conditions, which exhibit a range of QRS complex morphologies, noise levels, and motion artifacts. This work can be considered a cautionary tale about the impact of QRS detectors on HRV metrics errors, offering evidence to guide researchers and practitioners in selecting detection algorithms best suited to specific HRV features and recording conditions.

## II. Material and Methods

To evaluate the impact of QRS detection methods on HRV metrics, we computed interval tachograms from a set of ECG recordings using multiple detection algorithms. Then, we computed different HRV metrics for each interval tachogram. Finally, we conducted a statistical analysis to examine the differences. The source code used in this work is available on GitHub and archived in Zenodo [22]. Next, we describe each step in detail.

### A. ECG Database

We used the set of publicly available ECG signals in the Glasgow University Database (GUDB) [23]. GUDB contains ECG signals from 25 subjects recorded under five different conditions: sitting, walking, performing a math test, using a handbike, and jogging. Each ECG was recorded simultaneously with two independent setups: (1) an elastic chest strap with two electrodes connected to an amplifier and (2) a second pair of electrodes loosely connected to another amplifier (see [16] for more details about the recording setup). We identify these setups as “chest strap” and “loose cables,” respectively. The recording of both setups was synchronized. All ECG signals were sampled at 250 Hz. We selected this database because, unlike other ECG datasets, it includes a variety of recording conditions and setups with differing levels of noise and artifacts, making it particularly well-suited for evaluating the performance of QRS detection algorithms [16].

The GUDB includes annotations with the heartbeat locations, which we consider as ground truth. However, some recordings lack these annotations due to excessive noise— specifically, 2 recordings with the chest strap and 19 with loose cables, most of which correspond to the jogging condition [16]. Given the limited number of annotated signals in the loose cable jogging condition, we excluded this setup and condition combination from further analysis. Fig. 2 shows an example of an ECG segment for the jogging conditions for the loose cables setup (upper panel) and the chest strap setup (lower panel) recorded simultaneously. We can observe that the ECG signal recorded with the loose cable setup is noisier and lacks R-peak annotations.

**Fig. 2:**
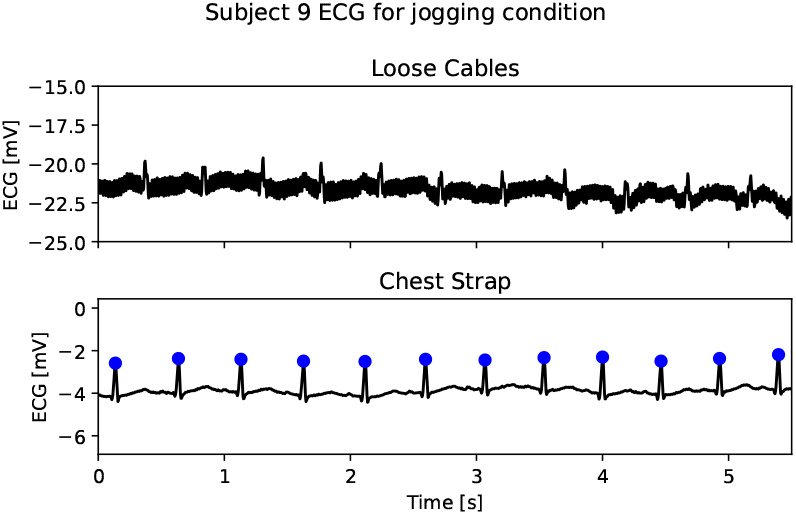
Example of raw ECG signal during the jogging condition. Upper: In black, the noisy ECG recording from the loose cables setup does not contain annotations for R-peaks. Lower: The same ECG recording, but now from the chest strap, including annotations for R-peaks.

### B. QRS detectors

We used eight QRS detection algorithms to obtain the interval tachogram from the ECG signals as implemented in the “ECG detectors” toolbox [24]. The following are the eight methods. In parentheses, we indicate the names used in the sequel, which follows the nomenclature used in the toolbox. Pan and Tompkins [25] (pan tompkins), Elgendi et al. [26] (two average), Kalidas and Tamil [27] (swt), Christov [28] (christov), Hamilton [29] (hamilton), Matched filter detector [24] (matched filter), Engelse and Zeelenberg [30] (engzee), and Zong et al. [31] (wqrs).

We excluded the Engzee detector in the chest strap setup because the detector could not detect most of the R-peaks (less than 5%) in seven cases (subject 11 while walking; subject 16 while sitting, using a handbike, and jogging; and subject 21 while sitting, performing a math test, and using a handbike). We illustrate this situation in Figure 3, where we show the ECG for subject 16 recorded using the chest strap during the sitting condition, the annotated peaks, the only peak detected by the Engzee detector, and, as a comparison, the peaks detected by the two average detector (the offsets between the R-peaks and the detections made by the two average method is normal, and it is not a problem as far as this offset is constant across the QRS complexes of the recording [16]).

**Fig. 3:**
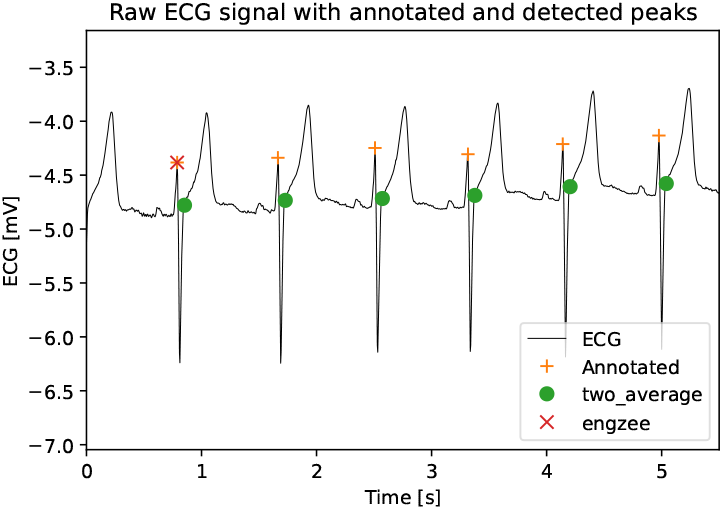
ECG signal of subject 16 during the sitting condition recorded with the chest strap setup. Orange plus signs mark the annotated peak, the red cross marks the only peak detected by Engzee, and the green dots mark the peaks detected by the two average method.

### C. Heart Rate Variability Metrics

We computed 23 HRV metrics for each interval tachogram obtained from the QRS detectors as well as for the corresponding tachogram based on annotated R-peaks. Table I lists and briefly describes the HRV metrics used in this study. The table also shows the analysis domain for each metric (time, frequency, or non-linear domain). See Pham et al. [32] for more details about how each metric is computed. We computed all the metrics using the Neurokit2 toolbox [33]. This set of 23 HRV metrics represents the ones commonly used in research and clinical applications [34].

**TABLE I:**
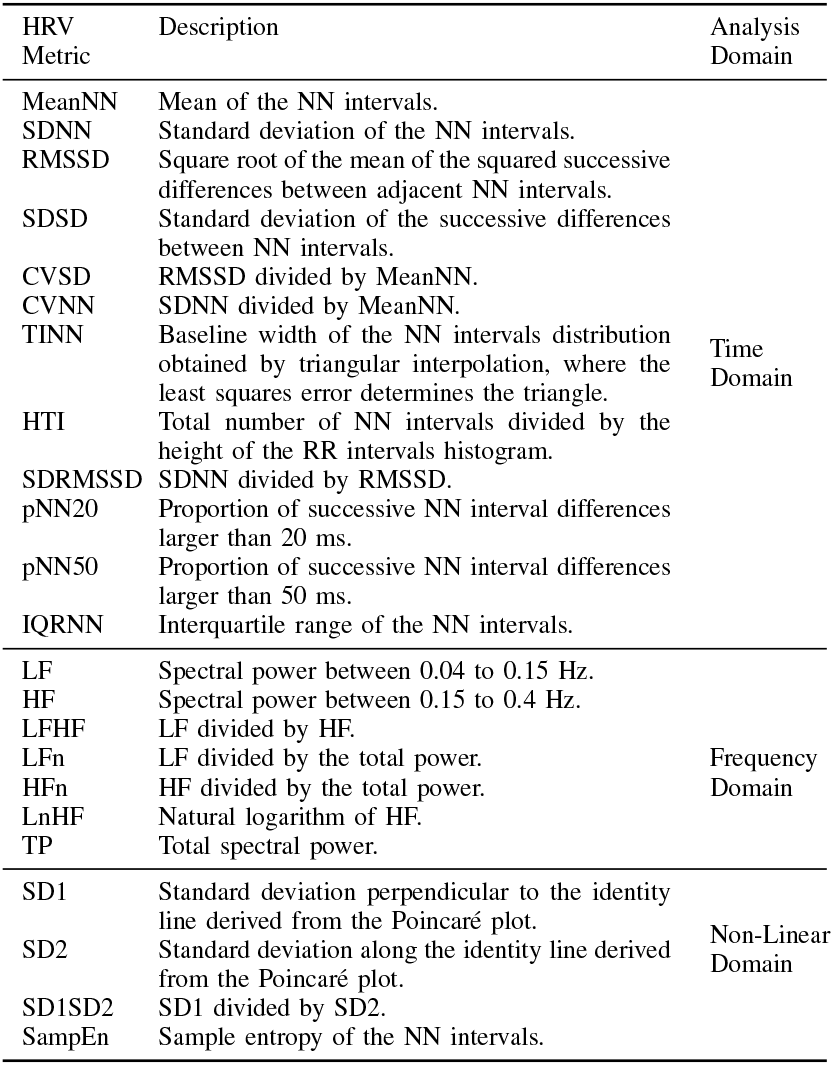
List of used HRV metrics.

### D. Concordance correlation coefficient

We used Lin’s concordance correlation coefficient (CCC) [35] to evaluate the agreement between the HRV metrics computed from annotated signals and the HRV metrics computed using the QRS detected by the different methods. As the comparison is made using the gold standard annotated data, the CCC can be interpreted as a measure of the accuracy of the QRS detector applied to the ECG signal [36]. Its values range from 0 to *±* 1, where 0 means no concordance and 1 means perfect concordance.

We computed the CCC for each pair of annotated data and the combination of HRV metrics, QRS detectors, and conditions. This analysis was conducted for the chest strap and loose cables setup.

For the chest strap setup, this analysis included seven QRS detectors (two average, matched filter, swt, engzee, christov, hamilton, and pan tompkins) and five experimental conditions (sitting, math, walking, handbike, and jogging). For the loose cables setup, the analysis included eight QRS detectors (two average, matched filter, swt, engzee, christov, hamilton, pan tompkins, and wqrs) and four conditions (sitting, math, walking, and handbike).

To help understand the meaning of the CCC, we show, for a given setup, HRV metric, and QRS detector, a scatter and regression plot. The plot includes one point per subject, with horizontal and vertical coordinates corresponding to the HRV metric value obtained using the annotated R-peak location and the R-peak location obtained from the QRS detector, respectively. The plot also includes the line obtained from a simple linear regression model where the independent and dependent variables are the HRV metric value obtained using the annotated R-peak location and the R-peak location obtained from the QRS detector, respectively. Each figure shows the scatter and regression plot for all conditions, with a different color for each condition. As a reference, we add a 45-degree angle dotted line representing where points with perfect matching between the annotated and measured HRV metric values lie.

### E. Performance evaluation and ranking of QRS Detectors

For each HRV metric, we evaluated the performance of every QRS detector based on rank and CCC. The rank reflects the relative position of each method within each condition, where a lower rank indicates better performance. This ranking was determined by the CCC values, where methods with higher CCC values received better ranks. CCC values larger than 0.8 were considered indicative of high-quality performance [37]. We have highlighted in bold all CCC values greater than 0.8 to help identify top-performing methods.

## III.Results

### A. Ranking of QRS detectors

We evaluated QRS detectors using both CCC values and rank to provide a comprehensive view of their performance, highlighting the most accurate and consistent methods across different experimental conditions. In this section, we show three examples of HRV metrics and the performance evaluation of the QRS detectors for every experimental condition. In the Supplementary Material section, we show the rank according to CCC values for every HRV metric and experimental condition.

As shown in Table II, when analyzing the MeanNN metric, the swt method consistently achieved high CCC values (above 0.99) and ranked in the top positions (1st or 2nd) for most conditions, demonstrating superior performance across all experimental scenarios. Similarly, the two average method performed well, particularly in the sitting and maths conditions, with CCC values surpassing 0.99 and ranking highly. Other methods, such as hamilton and matched filter, exhibited reasonable performance, but with lower CCC values in specific conditions like handbike and jogging, resulting in slightly lower ranks. The methods pan tompkins, christov, and wqrs showed more variability in performance, with a few CCC values above 0.8 but less consistency in their rankings.

**TABLE II:**
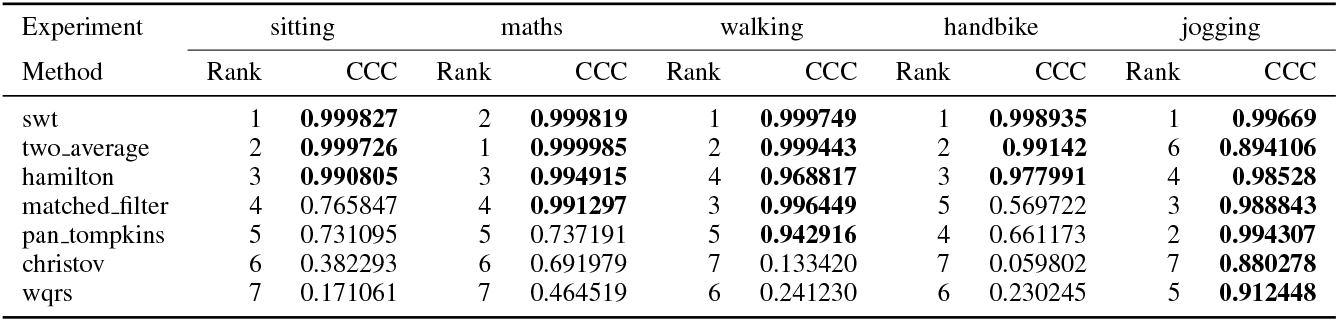
Ranking chest strap setup - Metric MeanNN.

In the analysis of the LFHF HRV metric (see Table III), the matched filter method was the top performer in the sitting and walking conditions, achieving the highest CCC values (above 0.9) in these conditions. It also ranked well in the maths condition (5th rank), but its CCC value in the handbike and jogging conditions (0.35) was much lower, reflecting reduced performance in these conditions. Similarly, christov demonstrated excellent performance in the maths, walking, and handbike conditions, with CCC values above 0.9, but its performance in jogging (rank 4, CCC 0.20) was significantly lower. The wqrs method showed consistent performance, with CCC values over 0.8 in the walking condition. However, its performance dropped in the sitting and maths conditions, ranking 3rd and 4th with lower CCC values of 0.80 and 0.66. The two average method performed well in the maths condition (rank 2, CCC 0.92) but showed weaker results in the walking, handbike, and sitting conditions, with CCC values below 0.8. The hamilton method consistently ranked in the lower positions for most conditions, with CCC values across all conditions mostly below 0.8, especially in the jogging condition, where the CCC value dropped drastically (0.09). Finally, swt and pan tompkins had the lowest CCC values across all experimental conditions, highlighting their relatively poor performance compared to the other methods.

**TABLE III:**
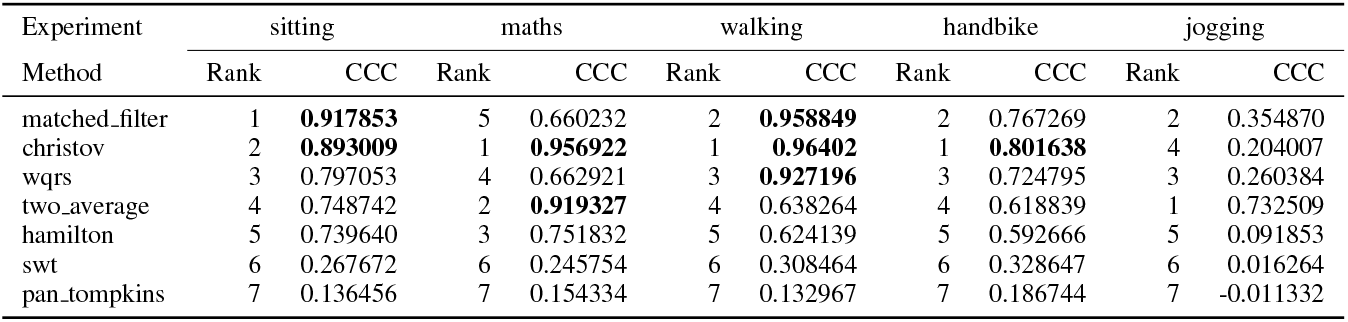
Ranking chest strap setup - Metric LFHF.

**TABLE IV:**
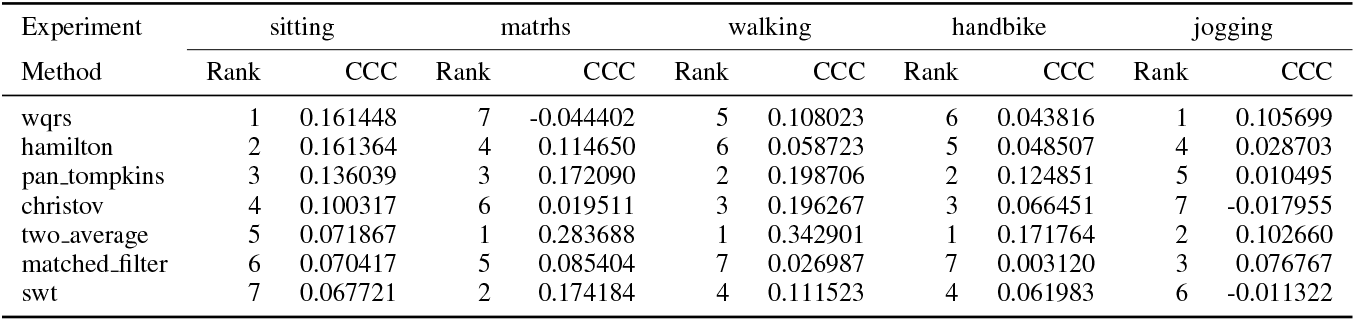
Ranking chest strap setup - Metric TINN.

In the case of the TINN HRV metric, we observed generally low performance from all QRS detection methods, as reflected by CCC values, which were all below 0.8 and mostly remained under 0.2. The two average method stands out with slightly better performance in certain conditions, but still falls short of achieving meaningful accuracy in most cases. The methods wqrs, hamilton, and christov show consistent poor performance across the conditions, with many CCC values being close to zero. Only a few CCC values exceed 0.2, indicating that the methods tested do not perform well for the TINN HRV metric.

In general, these analyses provide a comprehensive overview of the QRS detectors’ performance. CCC values and rankings reflect the varying levels of accuracy and consistency across the different experimental conditions, highlighting the varying effectiveness of the methods in detecting QRS complexes. Furthermore, it is shown how different HRV metrics are more or less affected by the performance of the QRS detectors.

While some methods, such as the swt and two average, consistently performed well for time-domain metrics like MeanNN and the chest strap setup, others, including wqrs, and christov, exhibited poor performance, with CCC values predominantly below 0.8. Overall, the analysis emphasizes the challenges in achieving high performance across all conditions and metrics, underscoring the need for further improvements in QRS detection methods.

To illustrate these differences more clearly, Figure 4 shows CCC values for three HRV metrics—MeanNN, TINN, and LFHF—using the chest strap setup across all QRS detection algorithms. The plot includes dotted reference lines indicating thresholds of concordance strength. For MeanNN, most detectors achieved moderately strong to very strong agreement, with CCC values frequently exceeding 0.8, indicating that this metric is relatively robust to detection variability in the chest strap configuration.

**Fig. 4:**
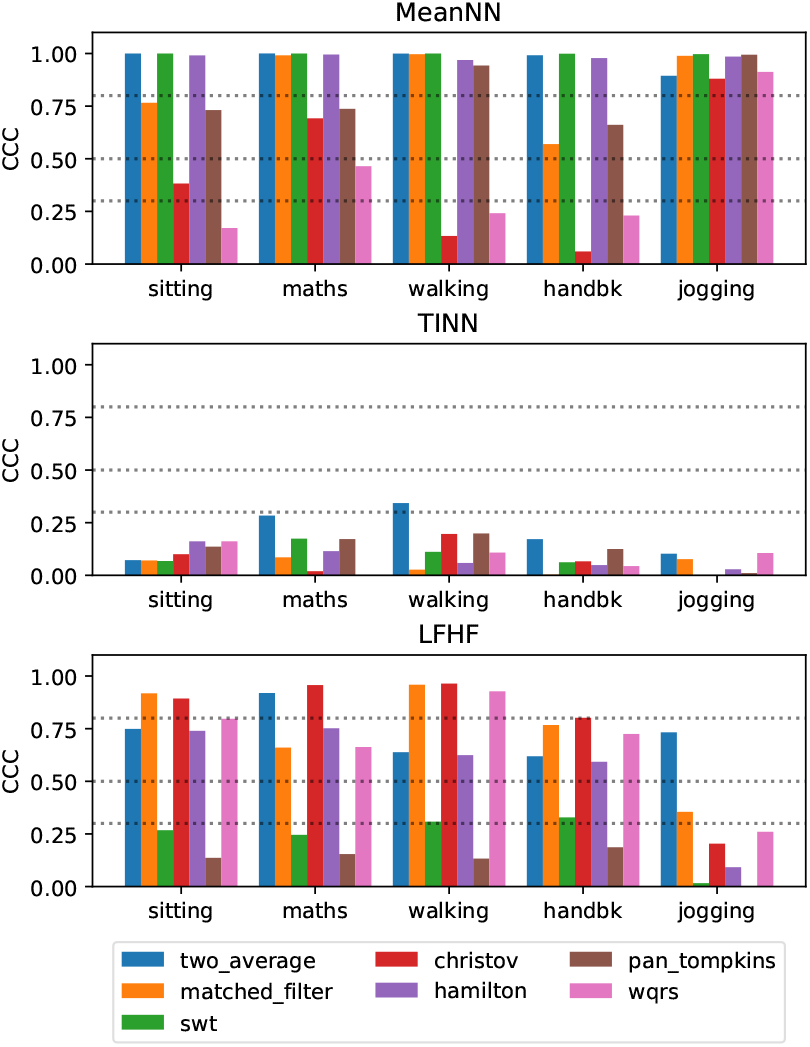
CCC for HRV metrics MeanNNN, TINN, and LFHF and the chest strap setup. The dotted lines show the concordance thresholds: “very strong” above 0.8, “moderately strong” between 0.5 and 0.8, “fair” between 0.3 and 0.5, and “poor” for less than 0.3.

In contrast, TINN exhibited consistently low concordance across all detectors, with CCC values predominantly falling in the “poor” range (below 0.3), indicating that this metric is highly sensitive to R-peak detection errors in this setup. On the other hand, the LFHF metric showed a mixed pattern, with CCC values ranging from “poor” to “moderately strong”, depending on the detector.

Figure 5 provides further insight into the agreement between annotated and detector-derived HRV values. It shows scatter plots for three HRV metrics (MeanNN, LFHF, and TINN), where each point corresponds to a subject’s measurement under various conditions, along with the associated linear regression lines and CCC values.

**Fig. 5:**
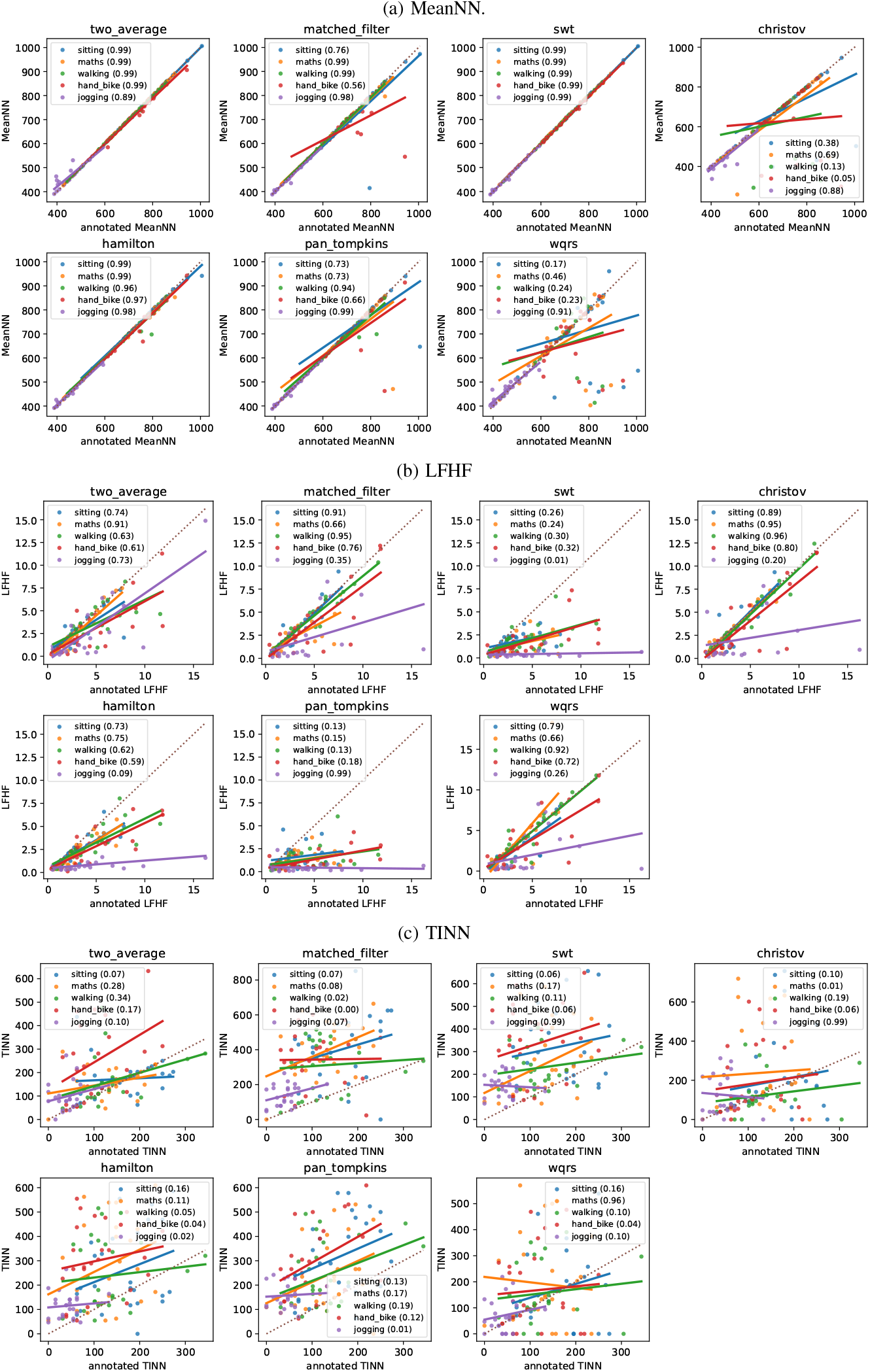
Interpretation of CCC. Each plot corresponds to the chest strap setup, a QRS detector, and HRV metrics (a) MeanNN, (b) LFHF, and (c) TINN. Each point represents a subject, with horizontal and vertical coordinates corresponding to HRV values from annotated and QRS detector-derived R-peak locations, respectively. A linear regression line is included, with annotated and detected HRV values as independent and dependent variables. Conditions are color-coded with CCC values indicated in the legend. A 45-degree dotted line indicates perfect agreement.

### B. HRV metrics ranking

Table V presents the percentage of CCC values greater than 0.8 for the HRV metrics analyzed in this article, for the chest strap and loose cables setup, which allows quantifying the performance of QRS detection methods. Notably, for the MeanNN metric, the loose cables setup demonstrated substantially better performance, with 84.4% of CCC values exceeding 0.8, in contrast to just 62.9% observed with the chest strap configuration. This suggests that the loose cables setup yields better results for detecting QRS complexes, which is then reflected in the computation of this metric. In contrast, the LF metric shows almost identical performance between the two setups, with 51.4% of CCC values above 0.8 for the chest strap setup and 53.1% for the loose cables setup, indicating a minimal difference in performance.

**TABLE V:** Comparison of CCC values for all HRV metrics between chest strap and loose cables setups. The values indicate the percentage of instances in which each metric achieved a CCC greater than 0.8, aggregated across all experimental conditions and QRS detection methods.

| Chest strap |  | Loose cables |  |
| --- | --- | --- | --- |
| Metric | CCC>0.8% | Metric | CCC>0.8% |
| MeanNN | 62.9 | MeanNN | 84.4 |
| LF | 51.4 | LF | 53.1 |
| HF<sub>n</sub> | 37.1 | IQRNN | 43.8 |
| pNN20 | 28.6 | pNN20 | 31.3 |
| IQRNN | 28.6 | SD2 | 28.1 |
| SD2 | 25.7 | HTI | 25 |
| HTI | 22.9 | HF<sub>n</sub> | 25 |
| LFHF | 22.9 | pNN50 | 18.8 |
| LnHF | 20 | LnHF | 15.6 |
| HF | 14.3 | LFHF | 12.5 |
| LF<sub>n</sub> | 11.4 | SampEn | 12.5 |
| TP | 11.4 | TP | 9.4 |
| SDNN | 8.6 | SDNN | 6.3 |
| CVNN | 8.6 | CVNN | 6.3 |
| SampEn | 5.7 | HF | 6.3 |
| pNN50 | 2.9 | SDRMSSD | 3.1 |
| RMSSD | 0 | LF<sub>n</sub> | 3.1 |
| SDSD | 0 | RMSSD | 0 |
| CVSD | 0 | SDSD | 0 |
| TINN | 0 | CVSD | 0 |
| SDRMSSD | 0 | TINN | 0 |
| SD1 | 0 | SD1 | 0 |
| SD1SD2 | 0 | SD1SD2 | 0 |

The HFn metric demonstrates a more notable disparity, with the chest strap setup performing better, as 37.1% of its CCC values exceed 0.8, while the loose cables setup only achieves 25%. This suggests that the chest strap setup may be more suitable for detecting QRS complexes using this particular metric. Similarly, for the HRV pNN20 metric, the loose cables setup again shows a slight edge with 31.3% of CCC values surpassing 0.8, compared to 28.6% in the chest strap setup.

For other metrics like IQRNN, SD2, and HTI, the performance tends to be lower, with the loose cables setup showing slightly better results, especially for IQRNN, which achieves 43.8% CCC values above 0.8 compared to 28.6% for the chest strap setup. However, several metrics, such as TINN, SD1, and SD1SD2, exhibit extremely low performance in both setups, with no CCC values exceeding 0.8. This indicates that the QRS detection methods do not perform well with these metrics under either setup.

In general, the loose cables setup provides superior performance across most HRV metrics, particularly for MeanNN where higher percentages of CCC values exceed 0.8. However, the chest strap setup performs better for certain metrics, such as HFn and LFHF, surpassing the loose cables setup in those cases. Despite these variations, both setups struggle significantly with metrics like TINN, SD1, SD1SD2, SDSD, CVSD, and RMSSD, where performance is consistently poor, highlighting the need for further refinement in QRS detection methods for these specific metrics.

## IV. Discussion

This study provides a comprehensive evaluation of how different QRS detection algorithms affect the accuracy of heart rate variability (HRV) metrics under realistic ECG acquisition conditions. Using real ECG data from 25 subjects recorded under various cognitive and physical activities, we examined the performance of eight widely used QRS detectors across two recording setups (chest strap and loose cables), highlighting their variable reliability depending on the HRV metric and signal quality.

Our results show that while some detectors perform better than others, none is the best under all conditions. This finding highlights the importance of carefully selecting a detector based on the conditions under which ECG signals are recorded, as well as the need for visual inspection of the detected R-peaks. The results also show that MeanNN exhibits the highest agreement with the ground truth. This is consistent with the fact that averaging RR intervals reduces the impact of potential errors introduced by the QRS detector. Conversely, the metrics showing the lowest agreement are those that more strongly amplify variability in RR intervals, such as SD1 and SD1SD2. These findings underscore the importance of considering ECG signal quality as a key criterion when selecting HRV metrics for a given study.

Recognizing the variability in QRS detector performance across different conditions, Porr and Macfarlane [16] proposed the JF metric as a unified measure of detection performance. It is a value between 0 and 100%, with 100% corresponding to a perfect match with respect to the annotated R-peaks. On the other hand, the CCC quantifies the agreement between an HRV metric computed from annotated R-peaks and the same metric derived from detected R-peaks, where a value of 1 indicates perfect agreement. We examined the relationship between JF and CCC across detectors and conditions. Each panel in Fig. 6 presents a scatter plot and linear regression for a specific HRV metric, with one point per detector-condition combination and separate regression lines for each setup. The *R*^2^ values for the chest strap and loose cables setup are in parentheses, respectively. A priori, a positive correlation between JF and CCC is expected: higher QRS detection accuracy should yield better HRV agreement. While this is true for some metrics and setups (e.g., pNN20 for both setups and LFHF for the chest strap setup), it does not hold for all cases (e.g., MeanNN and SD1 for both setups). These results highlight the complex interplay between R-peak detection accuracy and HRV computation, underscoring that some metrics are more sensitive to detection errors than others.

**Fig. 6:**
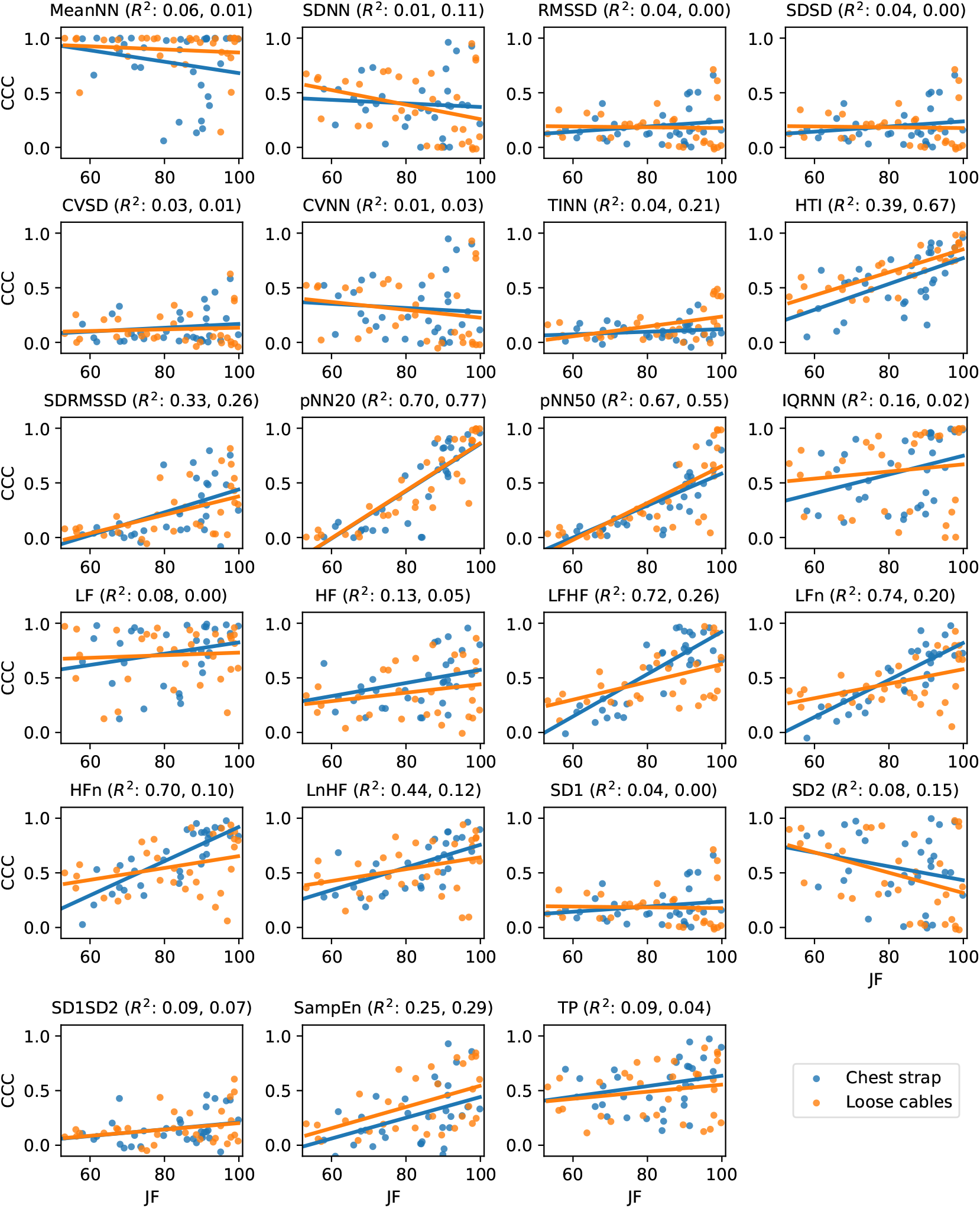
Correlation between JF and CCC. Each panel shows a scatter and regression plot for each HRV metric. The plot includes one point per condition and QRS detector. Blue and orange points correspond to the chest strap and loose cables setup, respectively. The lines represent a simple regression model for each setup. The corresponding *R*^2^ value for the chest strap and loose cables setup is shown in parentheses in the title.

Petelczyc et al. studied the effect of limited sampling frequencies and R-peaks detection errors in HRV metrics [18]. They modeled the interval tachogram as the sum of a ground truth time interval time series plus white Gaussian noise. Then, they characterized the error for different HRV metrics as a function of the noise level using Monte Carlo simulations. They found that MeanNN is the most robust metric, followed by SDNN and RMSSD, and that LF and LFHF are the least robust. Along the same lines, Rohr et al. [21] studied the effect of interval tachogram errors on HRV metrics. As ground truth, they used interval tachograms obtained from real restingstate ECG signals using a single QRS detector (pan tompkins). Then, they used synthetic noise and Monte Carlo simulations to determine how sensitive the different metrics are to this noise. They found that the most robust HRV metrics are MeanNN, LFHF ratio, and pNN50, and the least robust are LF and SDNN. These results are somewhat consistent with our findings, but we also observe some differences. Three reasons explain these differences. The first one is that these two studies did not evaluate the effect of QRS detectors. Instead, they considered the output of a given detector as ground truth. The second one is that they only considered ECG signals obtained under a single recording condition. The third one is that they used synthetic noise. In summary, and unlike previous works, our results highlight the effects of the QRS detector and, at the same time, consider a broad set of ECG signals representative of the recording conditions observed in practice.

A methodological limitation of our study is that ECG preprocessing was restricted to the steps embedded within each QRS detection algorithm’s implementation. Although, in principle, studying the effect of different preprocessing steps and their configurations could be possible, the search space is so large that this would render the problem infeasible. In addition, the goal of this work was not to identify a preprocessing pipeline to optimize the robustness of specific HRV metrics, but rather to highlight the impact of different QRS detection methods on HRV outcomes. Similarly, we did not evaluate the effect of interval tachogram correction methods [38], since introducing this additional factor to our study would significantly increase the complexity of the analysis, making the interpretation of the results more difficult. Future work will specifically address the influence of these correction methods.

Another limitation of our work is the number of subjects in the GUDB database. Although working with a larger number of subjects could make the results more statistically robust, the GUDB database is unique regarding its recording conditions. As noted by Porr et al. [16], it is the only annotated database that includes a range of noise levels and artifacts resembling those encountered in practical ECG recordings.

Doherty et al. underscore the critical need for robust validation of consumer wearable technologies, particularly as these devices are increasingly used in clinical and everyday health monitoring [39]. Their review revealed that HRV estimation from consumer wearables is generally valid only when recordings are taken at rest, with accuracy diminishing under movement or physical activity. Combined with our findings, this highlights the importance of examining how beat detection algorithms influence HRV metric errors. A study similar to ours, but focused specifically on the interaction between detection algorithms and wearable-derived HRV under varying conditions, would be a valuable direction for future research.

## V. Conclusion

Our results demonstrate that no single QRS detection algorithm consistently outperforms others across all conditions and HRV metrics. While detectors such as swt and two average generally performed well for time-domain metrics like MeanNN, frequency-domain, and non-linear metrics (e.g., LFHF) were significantly more sensitive to detection errors. Metrics such as TINN, SD1, and SD1SD2 consistently showed poor concordance across all detectors and setups. In our results, the loose cables ECG configuration often yielded higher CCC values than the chest strap setup, challenging the assumptions about its inferior signal quality. This finding suggests that different acquisition setups may be better suited for specific applications, depending on the context and the target HRV metrics. It is important to note that this study used real ECG recordings acquired under diverse experimental conditions, including rest, cognitive tasks, and physical activity. These data inherently have varying noise levels and different types of artifacts, providing a realistic benchmark for evaluating the performance of detectors in non-ideal conditions. This contrasts with prior studies such as those by [18], [21], which used synthetic models to simulate detection errors but did not account for algorithm-specific behavior under real-world variability. Our analysis also revealed that the relationship between QRS detection quality (measured using the JF metric) and HRV accuracy (measured using CCC) is metric-dependent and nonlinear, highlighting the need for context-specific evaluation. The choice of both the QRS detector and the ECG acquisition setup plays a critical role in determining the reliability of HRV analysis. This work provides practical guidance for clinicians and researchers in selecting QRS detection methods tailored to their analytical goals and recording conditions. Future research should investigate the impact of signal preprocessing and RR interval correction techniques to further improve the robustness of HRV metrics in noisy, reallife settings.

## Supplementary Material

### A. Chest strap setup Tables

**TABLE S1:**
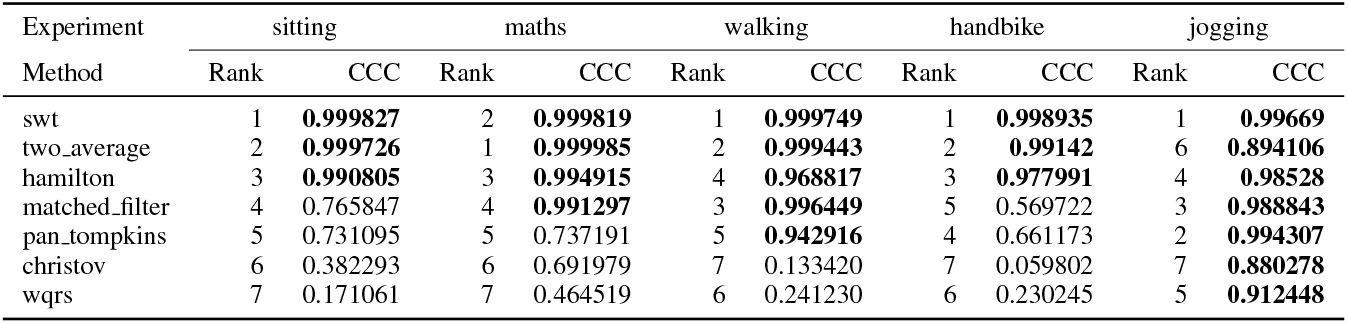
Ranking chest strap Setup - Metric MeanNN.

**TABLE S2:** Ranking chest strap Setup - Metric SDNN.

| Experiment | sitting |  | maths |  | walking |  | handbike |  | jogging |  |
| --- | --- | --- | --- | --- | --- | --- | --- | --- | --- | --- |
| Method | Rank | CCC | Rank | CCC | Rank | CCC | Rank | CCC | Rank | CCC |
| two_average | 1 | **0.926227** | 1 | **0.961532** | 1 | **0.882967** | 1 | 0.385674 | 6 | 0.358191 |
| swt | 2 | 0.730959 | 2 | 0.529891 | 2 | 0.705625 | 2 | 0.374442 | 1 | 0.678958 |
| hamilton | 3 | 0.718868 | 3 | 0.514997 | 6 | 0.333832 | 3 | 0.271945 | 4 | 0.448272 |
| pan_tompkins | 4 | 0.466793 | 4 | 0.399017 | 5 | 0.341470 | 4 | 0.205288 | 2 | 0.533700 |
| matched_filter | 5 | 0.392873 | 6 | 0.215985 | 4 | 0.383857 | 6 | 0.079377 | 3 | 0.520093 |
| christov | 6 | 0.375260 | 5 | 0.273330 | 3 | 0.399978 | 5 | 0.190409 | 5 | 0.408152 |
| wqrs | 7 | 0.001828 | 7 | 0.001902 | 7 | 0.012745 | 7 | 0.001166 | 7 | 0.030475 |

**TABLE S3:** Ranking chest strap Setup - Metric RMSSD.

| Experiment | sitting |  | maths |  | walking |  | handbike |  | jogging |  |
| --- | --- | --- | --- | --- | --- | --- | --- | --- | --- | --- |
| Method | Rank | CCC | Rank | CCC | Rank | CCC | Rank | CCC | Rank | CCC |
| two_average | 1 | 0.660828 | 1 | 0.504518 | 1 | 0.504472 | 1 | 0.147395 | 2 | 0.292810 |
| hamilton | 2 | 0.488923 | 2 | 0.353300 | 4 | 0.108979 | 2 | 0.127816 | 4 | 0.264515 |
| swt | 3 | 0.212442 | 4 | 0.141182 | 5 | 0.082958 | 4 | 0.091159 | 3 | 0.278898 |
| pan_tompkins | 4 | 0.179028 | 6 | 0.122813 | 6 | 0.056632 | 5 | 0.086786 | 6 | 0.165673 |
| christov | 5 | 0.173399 | 5 | 0.138462 | 3 | 0.120266 | 3 | 0.120002 | 1 | 0.401405 |
| matched_filter | 6 | 0.139544 | 3 | 0.158065 | 2 | 0.241002 | 6 | 0.055171 | 5 | 0.196088 |
| wqrs | 7 | 0.025871 | 7 | 0.003205 | 7 | 0.027151 | 7 | 0.026818 | 7 | 0.030919 |

**TABLE S4:**
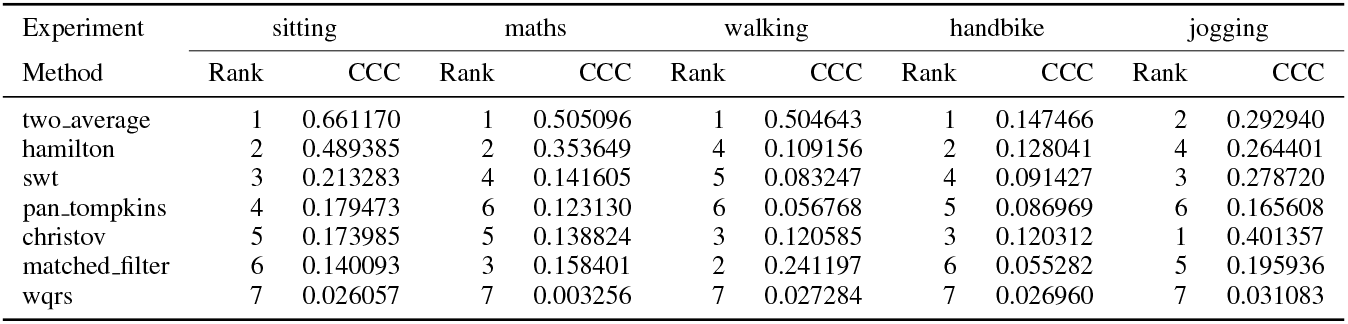
Ranking chest strap Setup - Metric SDSD.

**TABLE S5:**
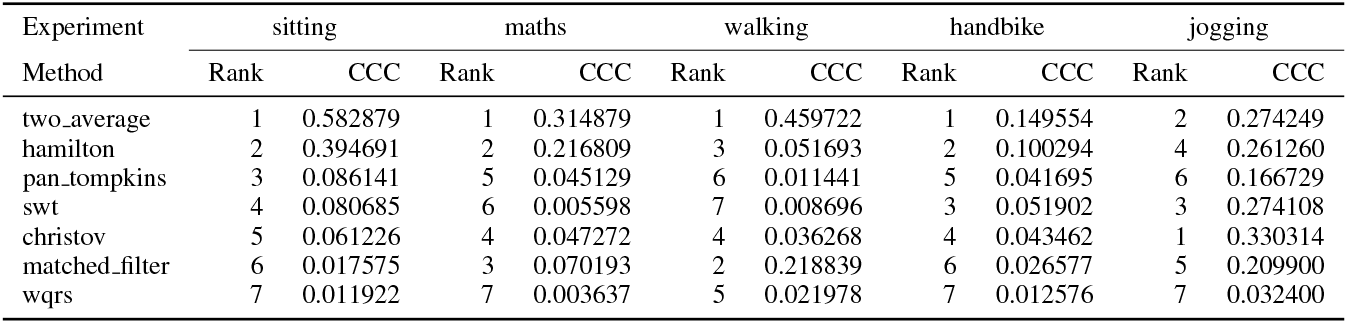
Ranking chest strap Setup - Metric CVSD.

**TABLE S6:** Ranking chest strap Setup - Metric CVNN.

| Experiment | sitting |  | maths |  | walking |  | handbike |  | jogging |  |
| --- | --- | --- | --- | --- | --- | --- | --- | --- | --- | --- |
| Method | Rank | CCC | Rank | CCC | Rank | CCC | Rank | CCC | Rank | CCC |
| two_average | 1 | **0.900111** | 1 | **0.947537** | 1 | **0.847531** | 1 | 0.371733 | 5 | 0.402802 |
| hamilton | 2 | 0.666470 | 2 | 0.435809 | 5 | 0.196984 | 3 | 0.232267 | 4 | 0.456893 |
| swt | 3 | 0.615692 | 3 | 0.426666 | 2 | 0.570674 | 2 | 0.342221 | 1 | 0.676308 |
| pan_tompkins | 4 | 0.228661 | 4 | 0.145104 | 4 | 0.198504 | 4 | 0.072727 | 3 | 0.521633 |
| matched_filter | 5 | 0.162436 | 5 | 0.114392 | 3 | 0.281359 | 6 | 0.026703 | 2 | 0.524708 |
| christov | 6 | 0.093651 | 6 | 0.096717 | 6 | 0.132037 | 5 | 0.029744 | 6 | 0.262742 |
| wqrs | 7 | -0.007734 | 7 | -0.000067 | 7 | 0.005626 | 7 | -0.010381 | 7 | 0.027768 |

**TABLE S7:**
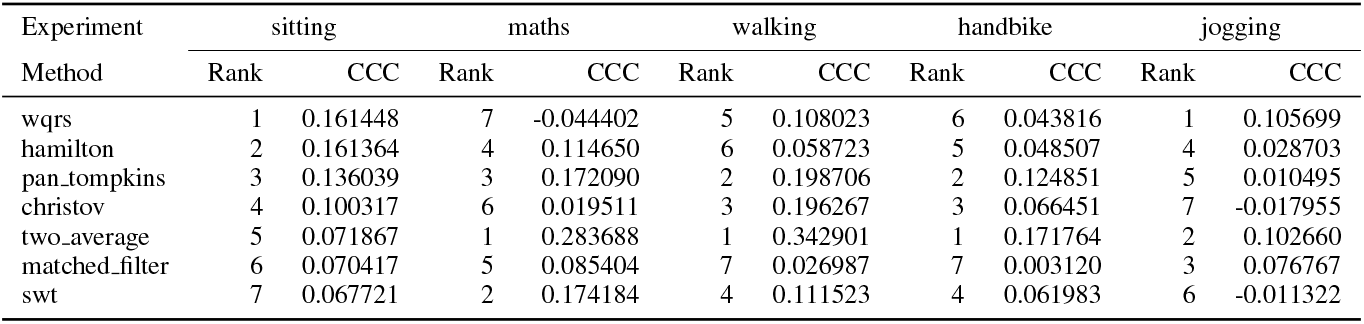
Ranking chest strap Setup - Metric TINN.

**TABLE S8:** Ranking chest strap Setup - Metric HTI.

| Experiment | sitting |  | maths |  | walking |  | handbike |  | jogging |  |
| --- | --- | --- | --- | --- | --- | --- | --- | --- | --- | --- |
| Method | Rank | CCC | Rank | CCC | Rank | CCC | Rank | CCC | Rank | CCC |
| two_average | 1 | **0.868249** | 2 | **0.911246** | 1 | **0.906041** | 1 | **0.819735** | 3 | 0.224334 |
| hamilton | 2 | **0.82115** | 3 | **0.868516** | 3 | 0.701456 | 2 | 0.757548 | 5 | 0.101725 |
| matched_filter | 3 | 0.633616 | 1 | **0.959345** | 2 | **0.804169** | 3 | 0.479738 | 2 | 0.371140 |
| christov | 4 | 0.572888 | 4 | 0.775743 | 6 | 0.465657 | 4 | 0.407839 | 4 | 0.178995 |
| swt | 5 | 0.566160 | 6 | 0.659525 | 5 | 0.543834 | 6 | 0.310830 | 7 | 0.046375 |
| pan_tompkins | 6 | 0.539740 | 5 | 0.668930 | 4 | 0.550540 | 7 | 0.309398 | 6 | 0.050119 |
| wqrs | 7 | 0.159122 | 7 | 0.544062 | 7 | 0.414817 | 5 | 0.354766 | 1 | 0.606458 |

**TABLE S9:**
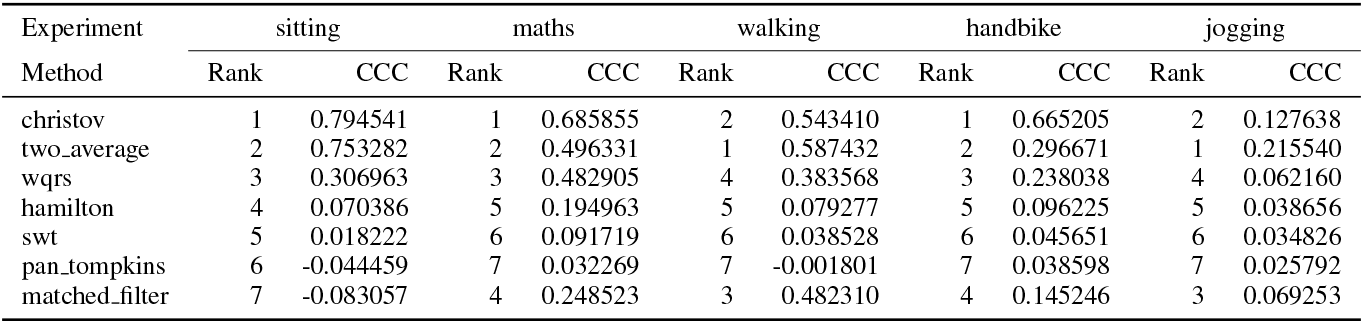
Ranking chest strap Setup - Metric SDRMSSD.

| Experiment | sitting |  | maths |  | walking |  | handbike |  | jogging |  |
| --- | --- | --- | --- | --- | --- | --- | --- | --- | --- | --- |
| Method | Rank | CCC | Rank | CCC | Rank | CCC | Rank | CCC | Rank | CCC |
| christov | 1 | 0.794541 | 1 | 0.685855 | 2 | 0.543410 | 1 | 0.665205 | 2 | 0.127638 |
| two_average | 2 | 0.753282 | 2 | 0.496331 | 1 | 0.587432 | 2 | 0.296671 | 1 | 0.215540 |
| wqrs | 3 | 0.306963 | 3 | 0.482905 | 4 | 0.383568 | 3 | 0.238038 | 4 | 0.062160 |
| hamilton | 4 | 0.070386 | 5 | 0.194963 | 5 | 0.079277 | 5 | 0.096225 | 5 | 0.038656 |
| swt | 5 | 0.018222 | 6 | 0.091719 | 6 | 0.038528 | 6 | 0.045651 | 6 | 0.034826 |
| pan_tompkins | 6 | -0.044459 | 7 | 0.032269 | 7 | -0.001801 | 7 | 0.038598 | 7 | 0.025792 |
| matched_filter | 7 | -0.083057 | 4 | 0.248523 | 3 | 0.482310 | 4 | 0.145246 | 3 | 0.069253 |

**TABLE S10:** Ranking chest strap Setup - Metric pNN20.

| Experiment | sitting |  | maths |  | walking |  | handbike |  | jogging |  |
| --- | --- | --- | --- | --- | --- | --- | --- | --- | --- | --- |
| Method | Rank | CCC | Rank | CCC | Rank | CCC | Rank | CCC | Rank | CCC |
| two_average | 1 | **0.853373** | 3 | **0.879934** | 3 | 0.724784 | 4 | 0.555486 | 4 | 0.001878 |
| christov | 2 | **0.823029** | 4 | **0.862642** | 4 | 0.620567 | 3 | 0.565210 | 2 | 0.058335 |
| wqrs | 3 | **0.821292** | 2 | **0.90586** | 2 | **0.816552** | 2 | 0.649311 | 1 | 0.066317 |
| matched_filter | 4 | **0.802114** | 1 | **0.955483** | 1 | **0.94249** | 1 | 0.673872 | 6 | 0.001449 |
| hamilton | 5 | 0.609350 | 5 | 0.584775 | 5 | 0.374181 | 5 | 0.127709 | 7 | -0.001147 |
| pan_tompkins | 6 | 0.258823 | 6 | 0.228802 | 6 | 0.078536 | 6 | 0.101714 | 3 | 0.003885 |
| swt | 7 | 0.063311 | 7 | 0.139638 | 7 | 0.069519 | 7 | 0.030017 | 5 | 0.001639 |

**TABLE S11:**
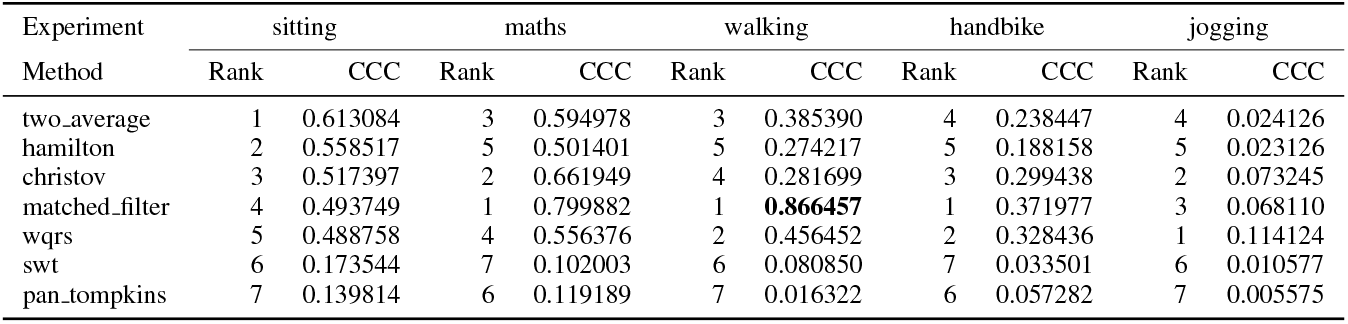
Ranking chest strap Setup - Metric pNN50.

**TABLE S12:** Ranking chest strap Setup - Metric IQRNN.

| Experiment | sitting |  | maths |  | walking |  | handbike |  | jogging |  |
| --- | --- | --- | --- | --- | --- | --- | --- | --- | --- | --- |
| Method | Rank | CCC | Rank | CCC | Rank | CCC | Rank | CCC | Rank | CCC |
| two_average | 1 | **0.966449** | 3 | **0.955239** | 2 | **0.896911** | 1 | **0.916979** | 6 | 0.206176 |
| hamilton | 2 | **0.951491** | 2 | **0.961193** | 3 | 0.786330 | 2 | 0.679264 | 2 | 0.621914 |
| swt | 3 | **0.898838** | 5 | 0.771171 | 4 | 0.760402 | 3 | 0.619800 | 4 | 0.291795 |
| matched_filter | 4 | 0.640193 | 1 | **0.985249** | 1 | **0.993367** | 4 | 0.214162 | 1 | **0.8852** |
| wqrs | 5 | 0.618089 | 4 | 0.774252 | 6 | 0.410512 | 7 | 0.164689 | 3 | 0.520334 |
| pan_tompkins | 6 | 0.368252 | 6 | 0.348346 | 5 | 0.460440 | 5 | 0.199626 | 5 | 0.243235 |
| christov | 7 | 0.336936 | 7 | 0.271336 | 7 | 0.295175 | 6 | 0.194178 | 7 | 0.137807 |

**TABLE S13:** Ranking chest strap Setup - Metric LF.

| Experiment | sitting |  | maths |  | walking |  | handbike |  | jogging |  |
| --- | --- | --- | --- | --- | --- | --- | --- | --- | --- | --- |
| Method | Rank | CCC | Rank | CCC | Rank | CCC | Rank | CCC | Rank | CCC |
| hamilton | 1 | **0.968898** | 4 | **0.966184** | 2 | **0.981996** | 1 | **0.894271** | 2 | 0.449038 |
| pan_tompkins | 2 | **0.931803** | 5 | **0.959703** | 6 | 0.636591 | 2 | **0.85968** | 1 | 0.647528 |
| swt | 3 | **0.928677** | 2 | **0.978349** | 5 | **0.815745** | 4 | 0.791655 | 3 | 0.395940 |
| two_average | 4 | **0.907307** | 1 | **0.99214** | 3 | **0.945452** | 3 | **0.831974** | 4 | 0.322172 |
| matched_filter | 5 | **0.838924** | 3 | **0.974084** | 1 | **0.98537** | 5 | 0.751358 | 5 | 0.264448 |
| christov | 6 | 0.732133 | 6 | 0.749597 | 4 | **0.841575** | 6 | 0.424430 | 7 | 0.124430 |
| wqrs | 7 | 0.547080 | 7 | 0.702918 | 7 | 0.625559 | 7 | 0.355985 | 6 | 0.215454 |

**TABLE S14:**
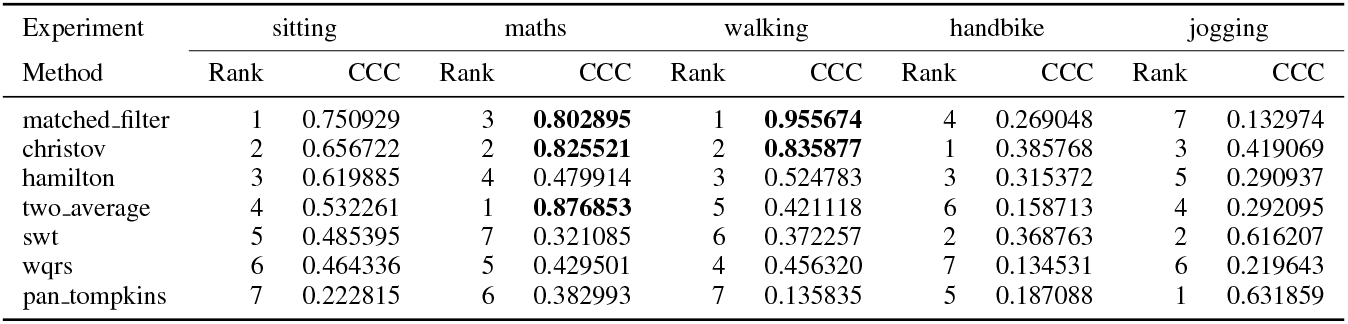
Ranking chest strap Setup - Metric HF.

**TABLE S15:**
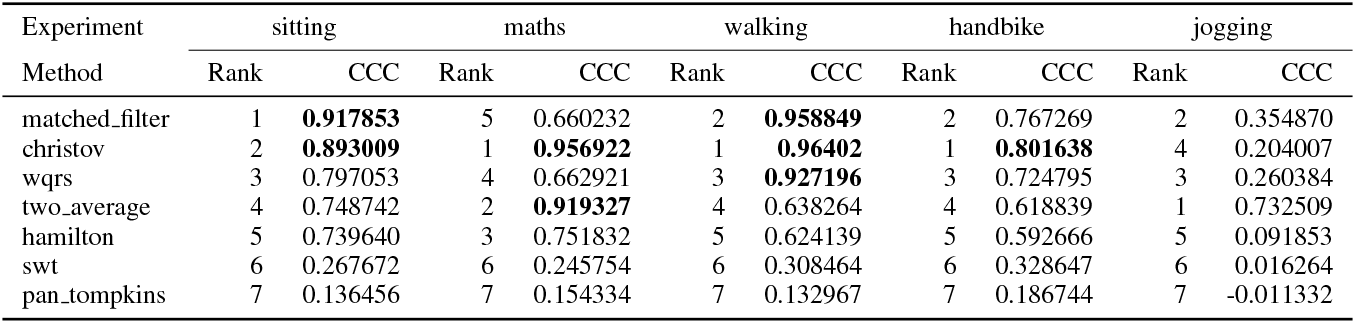
Ranking chest strap Setup - Metric LFHF.

**TABLE S16:** Ranking chest strap Setup - Metric LFn.

| Experiment | sitting |  | maths |  | walking |  | handbike |  | jogging |  |
| --- | --- | --- | --- | --- | --- | --- | --- | --- | --- | --- |
| Method | Rank | CCC | Rank | CCC | Rank | CCC | Rank | CCC | Rank | CCC |
| matched_filter | 1 | **0.893764** | 2 | 0.725491 | 1 | **0.977615** | 5 | 0.435677 | 2 | 0.376360 |
| christov | 2 | 0.729211 | 1 | 0.746784 | 2 | **0.932053** | 1 | 0.589243 | 4 | 0.285788 |
| hamilton | 3 | 0.717249 | 5 | 0.636385 | 4 | 0.741348 | 2 | 0.563787 | 5 | 0.101822 |
| two_average | 4 | 0.704785 | 3 | 0.700069 | 5 | 0.715492 | 3 | 0.492869 | 1 | 0.392200 |
| wqrs | 5 | 0.532282 | 4 | 0.685207 | 3 | **0.807244** | 4 | 0.476874 | 3 | 0.288355 |
| swt | 6 | 0.350199 | 7 | 0.234680 | 6 | 0.360481 | 6 | 0.307265 | 6 | -0.016441 |
| pan_tompkins | 7 | 0.178730 | 6 | 0.235514 | 7 | 0.162718 | 7 | 0.148078 | 7 | -0.051181 |

**TABLE S17:** Ranking chest strap Setup - Metric HFn.

| Experiment | sitting |  | maths |  | walking |  | handbike |  | jogging |  |
| --- | --- | --- | --- | --- | --- | --- | --- | --- | --- | --- |
| Method | Rank | CCC | Rank | CCC | Rank | CCC | Rank | CCC | Rank | CCC |
| matched_filter | 1 | **0.910672** | 3 | **0.836115** | 1 | **0.977206** | 4 | 0.579499 | 2 | 0.463139 |
| hamilton | 2 | **0.87016** | 5 | 0.769761 | 4 | **0.895724** | 1 | 0.696562 | 4 | 0.305044 |
| christov | 3 | **0.851454** | 2 | **0.858356** | 2 | **0.951721** | 3 | 0.618309 | 3 | 0.365844 |
| two_average | 4 | **0.839339** | 1 | **0.889217** | 3 | **0.91594** | 2 | 0.670003 | 1 | 0.514607 |
| wqrs | 5 | 0.648404 | 4 | **0.816754** | 5 | **0.869454** | 6 | 0.402590 | 5 | 0.292111 |
| swt | 6 | 0.627328 | 6 | 0.530164 | 6 | 0.567305 | 5 | 0.405927 | 6 | 0.128995 |
| pan_tompkins | 7 | 0.383173 | 7 | 0.517020 | 7 | 0.340651 | 7 | 0.205957 | 7 | 0.026879 |

**TABLE S18:** Ranking chest strap Setup - Metric LnHF.

| Experiment | sitting |  | maths |  | walking |  | handbike |  | jogging |  |
| --- | --- | --- | --- | --- | --- | --- | --- | --- | --- | --- |
| Method | Rank | CCC | Rank | CCC | Rank | CCC | Rank | CCC | Rank | CCC |
| matched_filter | 1 | **0.824671** | 1 | **0.896226** | 1 | **0.963778** | 5 | 0.371169 | 1 | 0.457736 |
| hamilton | 2 | 0.769182 | 3 | **0.874403** | 3 | 0.735755 | 1 | 0.590974 | 2 | 0.368428 |
| two_average | 3 | 0.696173 | 2 | **0.875787** | 2 | **0.815238** | 2 | 0.537665 | 3 | 0.360119 |
| wqrs | 4 | 0.666981 | 6 | 0.647450 | 5 | 0.503879 | 4 | 0.380608 | 5 | 0.314911 |
| christov | 5 | 0.618251 | 4 | **0.808401** | 4 | 0.630996 | 6 | 0.358069 | 7 | 0.272000 |
| swt | 6 | 0.503587 | 5 | 0.684875 | 6 | 0.442877 | 3 | 0.491059 | 4 | 0.337294 |
| pan_tompkins | 7 | 0.285825 | 7 | 0.583003 | 7 | 0.188540 | 7 | 0.295122 | 6 | 0.277731 |

**TABLE S19:**
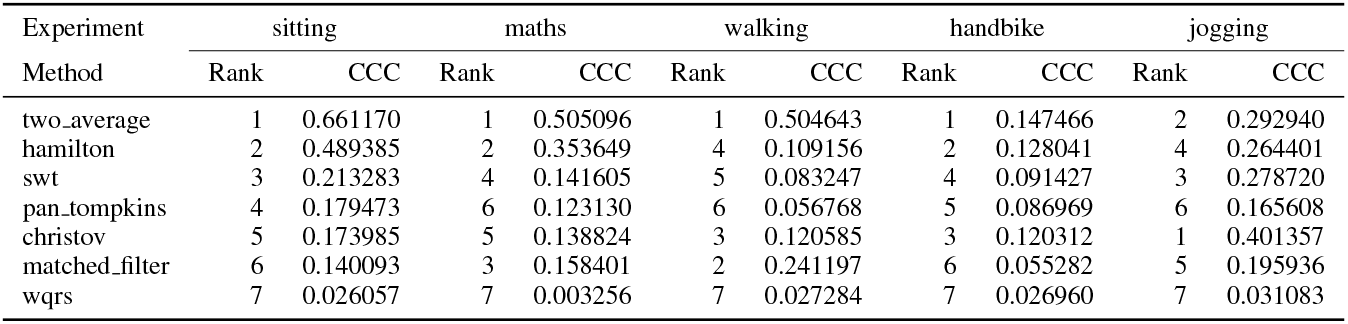
Ranking chest strap Setup - Metric SD1.

**TABLE S20:** Ranking chest strap Setup - Metric SD2.

| Experiment | sitting |  | maths |  | walking |  | handbike |  | jogging |  |
| --- | --- | --- | --- | --- | --- | --- | --- | --- | --- | --- |
| Method | Rank | CCC | Rank | CCC | Rank | CCC | Rank | CCC | Rank | CCC |
| swt | 1 | **0.973083** | 3 | **0.863095** | 2 | **0.941378** | 1 | 0.715176 | 1 | **0.890299** |
| two_average | 2 | **0.966667** | 1 | **0.996467** | 1 | **0.941719** | 2 | 0.556967 | 5 | 0.477484 |
| pan_tompkins | 3 | 0.794663 | 2 | **0.866788** | 4 | 0.517509 | 4 | 0.370589 | 2 | **0.83851** |
| hamilton | 4 | 0.791228 | 4 | 0.611326 | 6 | 0.415139 | 3 | 0.395720 | 4 | 0.652815 |
| christov | 5 | 0.653241 | 5 | 0.467332 | 3 | 0.688175 | 5 | 0.309017 | 6 | 0.429998 |
| matched_filter | 6 | 0.505689 | 6 | 0.294611 | 5 | 0.502601 | 6 | 0.106662 | 3 | 0.766426 |
| wqrs | 7 | -0.005559 | 7 | 0.021615 | 7 | 0.029716 | 7 | 0.008013 | 7 | 0.076466 |

**TABLE S21:** Ranking chest strap Setup - Metric SD1SD2.

| Experiment | sitting |  | maths |  | walking |  | handbike |  | jogging |  |
| --- | --- | --- | --- | --- | --- | --- | --- | --- | --- | --- |
| Method | Rank | CCC | Rank | CCC | Rank | CCC | Rank | CCC | Rank | CCC |
| two_average | 1 | 0.413381 | 1 | 0.459264 | 1 | 0.428001 | 3 | 0.121304 | 2 | 0.234195 |
| hamilton | 2 | 0.119804 | 2 | 0.351230 | 4 | 0.053167 | 1 | 0.153858 | 4 | 0.202872 |
| wqrs | 3 | 0.070010 | 4 | 0.136009 | 3 | 0.081437 | 4 | 0.115508 | 6 | 0.131620 |
| christov | 4 | 0.065346 | 7 | 0.058586 | 5 | 0.022356 | 5 | 0.088434 | 1 | 0.428107 |
| pan_tompkins | 5 | -0.012660 | 5 | 0.089665 | 7 | -0.025606 | 6 | 0.053535 | 7 | 0.107257 |
| swt | 6 | -0.013889 | 6 | 0.080585 | 6 | 0.008980 | 7 | 0.049387 | 5 | 0.176613 |
| matched_filter | 7 | -0.061495 | 3 | 0.231630 | 2 | 0.406334 | 2 | 0.123524 | 3 | 0.206139 |

**TABLE S22:**
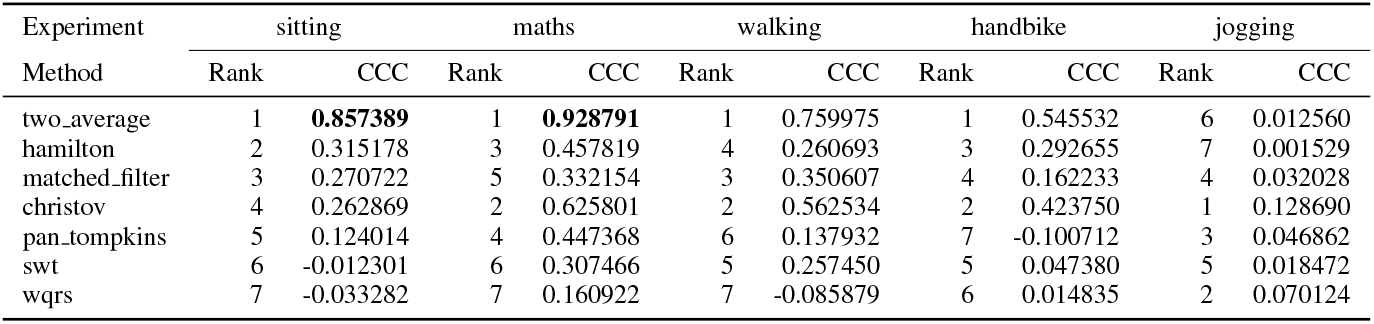
Ranking chest strap Setup - Metric HRV SampEn.

| Experiment | sitting |  | maths |  | walking |  | handbike |  | jogging |  |
| --- | --- | --- | --- | --- | --- | --- | --- | --- | --- | --- |
| Method | Rank | CCC | Rank | CCC | Rank | CCC | Rank | CCC | Rank | CCC |
| two_average | 1 | **0.857389** | 1 | **0.928791** | 1 | 0.759975 | 1 | 0.545532 | 6 | 0.012560 |
| hamilton | 2 | 0.315178 | 3 | 0.457819 | 4 | 0.260693 | 3 | 0.292655 | 7 | 0.001529 |
| matched_filter | 3 | 0.270722 | 5 | 0.332154 | 3 | 0.350607 | 4 | 0.162233 | 4 | 0.032028 |
| christov | 4 | 0.262869 | 2 | 0.625801 | 2 | 0.562534 | 2 | 0.423750 | 1 | 0.128690 |
| pan_tompkins | 5 | 0.124014 | 4 | 0.447368 | 6 | 0.137932 | 7 | -0.100712 | 3 | 0.046862 |
| swt | 6 | -0.012301 | 6 | 0.307466 | 5 | 0.257450 | 5 | 0.047380 | 5 | 0.018472 |
| wqrs | 7 | -0.033282 | 7 | 0.160922 | 7 | -0.085879 | 6 | 0.014835 | 2 | 0.070124 |

**TABLE S23:** Ranking chest strap Setup - Metric HRV TP.

| Experiment | sitting |  | maths |  | walking |  | handbike |  | jogging |  |
| --- | --- | --- | --- | --- | --- | --- | --- | --- | --- | --- |
| Method | Rank | CCC | Rank | CCC | Rank | CCC | Rank | CCC | Rank | CCC |
| matched_filter | 1 | 0.799702 | 2 | **0.896267** | 1 | **0.973079** | 4 | 0.366074 | 7 | 0.219747 |
| hamilton | 2 | 0.723000 | 4 | 0.718021 | 3 | 0.622488 | 3 | 0.434722 | 4 | 0.353986 |
| christov | 3 | 0.702092 | 3 | 0.757514 | 2 | **0.80771** | 5 | 0.242584 | 3 | 0.371112 |
| swt | 4 | 0.690009 | 6 | 0.610248 | 5 | 0.541975 | 1 | 0.498909 | 2 | 0.578033 |
| two_average | 5 | 0.675287 | 1 | **0.945045** | 4 | 0.616482 | 6 | 0.178573 | 5 | 0.328060 |
| wqrs | 6 | 0.553010 | 7 | 0.538984 | 6 | 0.425898 | 7 | 0.134087 | 6 | 0.250947 |
| pan_tompkins | 7 | 0.443180 | 5 | 0.621897 | 7 | 0.260981 | 2 | 0.441521 | 1 | 0.695478 |

### B. Loose cables setup Tables

**TABLE S24:** Ranking loose cables setup - Metric MeanNN.

| Experiment | sitting |  | maths |  | walking |  | handbike |  |
| --- | --- | --- | --- | --- | --- | --- | --- | --- |
| Method | Rank | CCC | Rank | CCC | Rank | CCC | Rank | CCC |
| two_average | 1 | **0.999756** | 1 | **0.999854** | 2 | **0.998426** | 1 | **0.995378** |
| pan_tompkins | 2 | **0.999478** | 2 | **0.999784** | 7 | 0.500420 | 4 | **0.982951** |
| swt | 3 | **0.999158** | 4 | **0.999417** | 1 | **0.999035** | 2 | **0.991009** |
| engzee | 4 | **0.995761** | 8 | 0.503876 | 8 | 0.140426 | 7 | **0.860273** |
| hamilton | 5 | **0.983377** | 6 | **0.993314** | 3 | **0.910407** | 3 | **0.984723** |
| matched_filter | 6 | **0.954152** | 5 | **0.995977** | 5 | **0.832857** | 5 | **0.981255** |
| christov | 7 | **0.871046** | 3 | **0.999693** | 4 | **0.838127** | 8 | 0.760028 |
| wqrs | 8 | **0.819758** | 7 | **0.96762** | 6 | 0.758387 | 6 | **0.959791** |

**TABLE S25:** Ranking loose cables setup - Metric SDNN.

| Experiment | sitting |  | maths |  | walking |  | handbike |  |
| --- | --- | --- | --- | --- | --- | --- | --- | --- |
| Method | Rank | CCC | Rank | CCC | Rank | CCC | Rank | CCC |
| two_average | 1 | **0.950109** | 1 | **0.819286** | 1 | 0.677455 | 5 | 0.458229 |
| pan_tompkins | 2 | 0.701269 | 3 | 0.645804 | 4 | 0.260782 | 3 | 0.604940 |
| swt | 3 | 0.693148 | 4 | 0.640718 | 2 | 0.672823 | 1 | 0.622480 |
| hamilton | 4 | 0.440654 | 5 | 0.267933 | 6 | 0.195157 | 4 | 0.577830 |
| matched_filter | 5 | 0.169097 | 6 | 0.002233 | 5 | 0.199393 | 6 | 0.317508 |
| christov | 6 | 0.163903 | 2 | 0.798179 | 3 | 0.337459 | 2 | 0.606103 |
| engzee | 7 | 0.097369 | 8 | -0.017851 | 7 | 0.030308 | 7 | 0.052268 |
| wqrs | 8 | -0.002496 | 7 | -0.013612 | 8 | 0.013230 | 8 | -0.000343 |

**TABLE S26:**
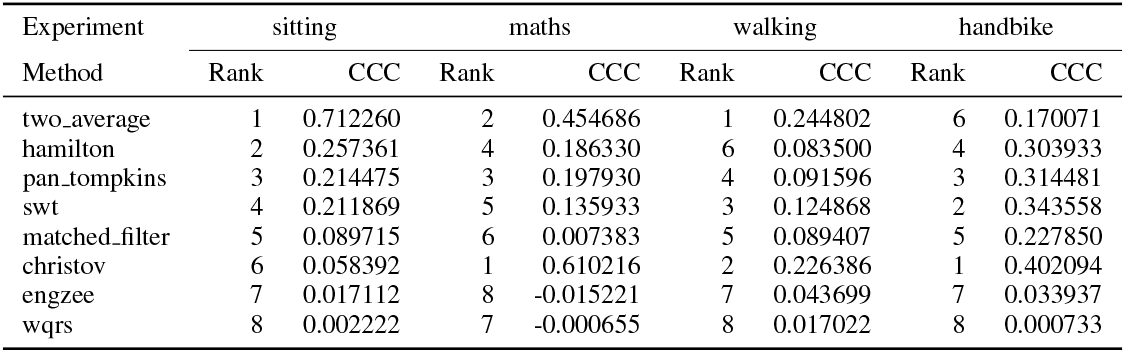
Ranking loose cables setup - Metric RMSSD.

| Experiment | sitting |  | maths |  | walking |  | handbike |  |
| --- | --- | --- | --- | --- | --- | --- | --- | --- |
| Method | Rank | CCC | Rank | CCC | Rank | CCC | Rank | CCC |
| two_average | 1 | 0.712260 | 2 | 0.454686 | 1 | 0.244802 | 6 | 0.170071 |
| hamilton | 2 | 0.257361 | 4 | 0.186330 | 6 | 0.083500 | 4 | 0.303933 |
| pan_tompkins | 3 | 0.214475 | 3 | 0.197930 | 4 | 0.091596 | 3 | 0.314481 |
| swt | 4 | 0.211869 | 5 | 0.135933 | 3 | 0.124868 | 2 | 0.343558 |
| matched_filter | 5 | 0.089715 | 6 | 0.007383 | 5 | 0.089407 | 5 | 0.227850 |
| christov | 6 | 0.058392 | 1 | 0.610216 | 2 | 0.226386 | 1 | 0.402094 |
| engzee | 7 | 0.017112 | 8 | -0.015221 | 7 | 0.043699 | 7 | 0.033937 |
| wqrs | 8 | 0.002222 | 7 | -0.000655 | 8 | 0.017022 | 8 | 0.000733 |

**TABLE S27:** Ranking loose cables setup - Metric SDSD.

| Experiment | sitting |  | maths |  | walking |  | handbike |  |
| --- | --- | --- | --- | --- | --- | --- | --- | --- |
| Method | Rank | CCC | Rank | CCC | Rank | CCC | Rank | CCC |
| two_average | 1 | 0.712546 | 2 | 0.454816 | 1 | 0.244821 | 6 | 0.170212 |
| hamilton | 2 | 0.257936 | 4 | 0.186665 | 6 | 0.083640 | 4 | 0.304300 |
| pan_tompkins | 3 | 0.215053 | 3 | 0.198372 | 4 | 0.091818 | 3 | 0.314883 |
| swt | 4 | 0.212504 | 5 | 0.136274 | 3 | 0.125034 | 2 | 0.343913 |
| matched_filter | 5 | 0.089855 | 6 | 0.007531 | 5 | 0.089439 | 5 | 0.228020 |
| christov | 6 | 0.058840 | 1 | 0.611701 | 2 | 0.226433 | 1 | 0.402222 |
| engzee | 7 | 0.017490 | 8 | -0.015179 | 7 | 0.043603 | 7 | 0.033919 |
| wqrs | 8 | 0.002267 | 7 | -0.000645 | 8 | 0.017081 | 8 | 0.000796 |

**TABLE S28:** Ranking loose cables setup - Metric CVSD.

| Experiment | sitting |  | maths |  | walking |  | handbike |  |
| --- | --- | --- | --- | --- | --- | --- | --- | --- |
| Method | Rank | CCC | Rank | CCC | Rank | CCC | Rank | CCC |
| two_average | 1 | 0.626233 | 2 | 0.379518 | 1 | 0.256359 | 6 | 0.131843 |
| hamilton | 2 | 0.130452 | 4 | 0.037948 | 6 | 0.050020 | 3 | 0.212430 |
| pan_tompkins | 3 | 0.076180 | 3 | 0.050528 | 7 | 0.029964 | 4 | 0.208142 |
| swt | 4 | 0.057979 | 5 | -0.000454 | 3 | 0.080696 | 2 | 0.253273 |
| matched_filter | 5 | 0.051855 | 8 | -0.037705 | 4 | 0.077461 | 5 | 0.170376 |
| wqrs | 6 | 0.005488 | 6 | -0.009857 | 8 | 0.018917 | 8 | 0.002814 |
| christov | 7 | -0.004892 | 1 | 0.404040 | 2 | 0.234545 | 1 | 0.339640 |
| engzee | 8 | -0.041020 | 7 | -0.022303 | 5 | 0.063818 | 7 | 0.042031 |

**TABLE S29:** Ranking loose cables setup - Metric CVNN.

| Experiment | sitting |  | maths |  | walking |  | handbike |  |
| --- | --- | --- | --- | --- | --- | --- | --- | --- |
| Method | Rank | CCC | Rank | CCC | Rank | CCC | Rank | CCC |
| two_average | 1 | **0.928812** | 1 | **0.813176** | 1 | 0.608417 | 4 | 0.427225 |
| pan_tompkins | 2 | 0.576134 | 4 | 0.516703 | 7 | 0.021562 | 2 | 0.499799 |
| swt | 3 | 0.536563 | 3 | 0.518810 | 2 | 0.522294 | 1 | 0.504920 |
| hamilton | 4 | 0.286725 | 5 | 0.152165 | 6 | 0.033302 | 3 | 0.480405 |
| matched_filter | 5 | 0.092375 | 8 | -0.053698 | 4 | 0.110472 | 6 | 0.249220 |
| wqrs | 6 | 0.001977 | 7 | -0.022294 | 8 | 0.011052 | 8 | -0.004556 |
| christov | 7 | -0.002448 | 2 | 0.769397 | 3 | 0.226914 | 5 | 0.344511 |
| engzee | 8 | -0.021047 | 6 | -0.017316 | 5 | 0.048034 | 7 | 0.076983 |

**TABLE S30:**
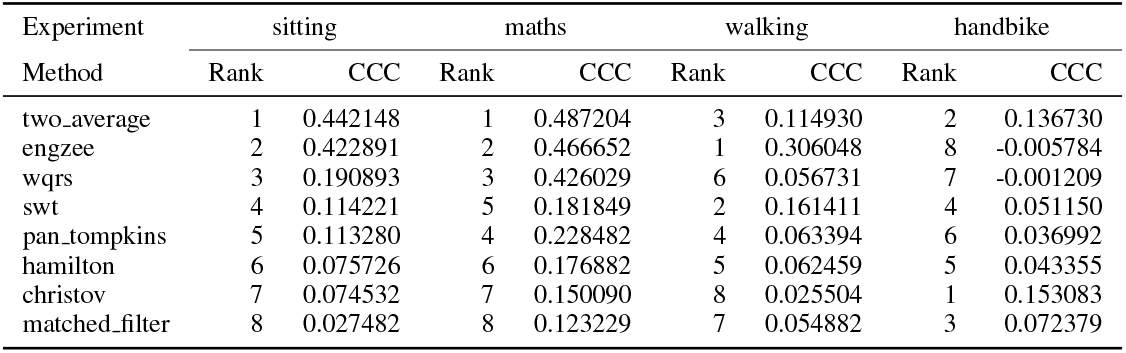
Ranking loose cables setup - Metric TINN.

| Experiment | sitting |  | maths |  | walking |  | handbike |  |
| --- | --- | --- | --- | --- | --- | --- | --- | --- |
| Method | Rank | CCC | Rank | CCC | Rank | CCC | Rank | CCC |
| two_average | 1 | 0.442148 | 1 | 0.487204 | 3 | 0.114930 | 2 | 0.136730 |
| engzee | 2 | 0.422891 | 2 | 0.466652 | 1 | 0.306048 | 8 | -0.005784 |
| wqrs | 3 | 0.190893 | 3 | 0.426029 | 6 | 0.056731 | 7 | -0.001209 |
| swt | 4 | 0.114221 | 5 | 0.181849 | 2 | 0.161411 | 4 | 0.051150 |
| pan_tompkins | 5 | 0.113280 | 4 | 0.228482 | 4 | 0.063394 | 6 | 0.036992 |
| hamilton | 6 | 0.075726 | 6 | 0.176882 | 5 | 0.062459 | 5 | 0.043355 |
| christov | 7 | 0.074532 | 7 | 0.150090 | 8 | 0.025504 | 1 | 0.153083 |
| matched_filter | 8 | 0.027482 | 8 | 0.123229 | 7 | 0.054882 | 3 | 0.072379 |

**TABLE S31:** Ranking loose cables setup - Metric HTI.

| Experiment | sitting |  | maths |  | walking |  | handbike |  |
| --- | --- | --- | --- | --- | --- | --- | --- | --- |
| Method | Rank | CCC | Rank | CCC | Rank | CCC | Rank | CCC |
| engzee | 1 | **0.991225** | 1 | **0.981772** | 1 | 0.731583 | 2 | 0.739309 |
| two_average | 2 | **0.864282** | 2 | **0.91259** | 4 | 0.612736 | 1 | 0.769752 |
| matched_filter | 3 | **0.843336** | 5 | **0.889301** | 8 | 0.383429 | 7 | 0.405639 |
| wqrs | 4 | 0.750240 | 4 | **0.89714** | 3 | 0.613405 | 3 | 0.657931 |
| christov | 5 | 0.744613 | 3 | **0.912458** | 2 | 0.644645 | 4 | 0.645029 |
| hamilton | 6 | 0.735596 | 7 | 0.703617 | 5 | 0.593991 | 6 | 0.482750 |
| pan_tompkins | 7 | 0.484595 | 6 | 0.723947 | 6 | 0.510323 | 5 | 0.495964 |
| swt | 8 | 0.391925 | 8 | 0.595161 | 7 | 0.417444 | 8 | 0.273143 |

**TABLE S32:**
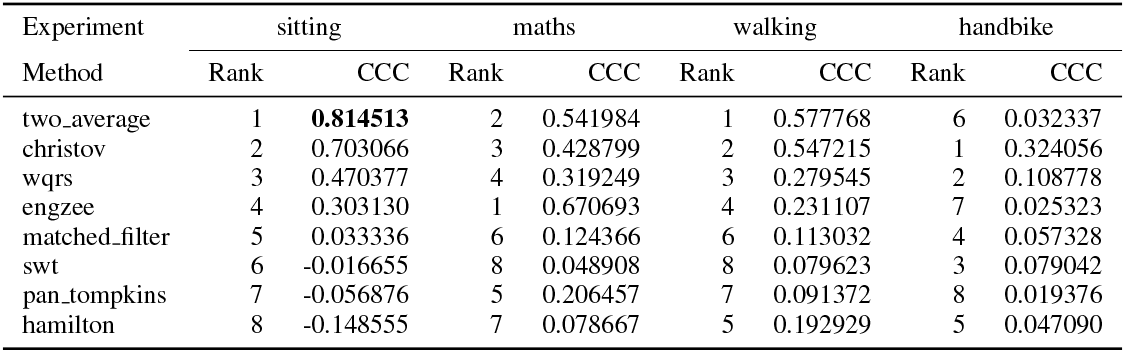
Ranking loose cables setup - Metric SDRMSSD.

**TABLE S33:** Ranking loose cables setup - Metric pNN20.

| Experiment | sitting |  | maths |  | walking |  | handbike |  |
| --- | --- | --- | --- | --- | --- | --- | --- | --- |
| Method | Rank | CCC | Rank | CCC | Rank | CCC | Rank | CCC |
| engzee | 1 | **0.995687** | 5 | **0.886109** | 3 | 0.516682 | 4 | 0.312148 |
| wqrs | 2 | **0.983206** | 2 | **0.994765** | 1 | 0.686669 | 1 | 0.712205 |
| christov | 3 | **0.900835** | 1 | **0.995087** | 2 | 0.525328 | 3 | 0.324116 |
| matched_filter | 4 | **0.898542** | 4 | **0.90416** | 5 | 0.284375 | 5 | 0.274587 |
| two_average | 5 | **0.891745** | 3 | **0.973361** | 4 | 0.439475 | 2 | 0.581241 |
| hamilton | 6 | 0.273484 | 7 | 0.229662 | 6 | 0.035836 | 7 | 0.004557 |
| pan_tompkins | 7 | 0.204195 | 6 | 0.364203 | 7 | 0.023831 | 6 | 0.027278 |
| swt | 8 | 0.200607 | 8 | 0.069095 | 8 | 0.004112 | 8 | -0.001023 |

**TABLE S34:** Ranking loose cables setup - Metric pNN50.

| Experiment | sitting |  | maths |  | walking |  | handbike |  |
| --- | --- | --- | --- | --- | --- | --- | --- | --- |
| Method | Rank | CCC | Rank | CCC | Rank | CCC | Rank | CCC |
| engzee | 1 | **0.984837** | 5 | 0.561615 | 3 | 0.189106 | 5 | 0.042294 |
| wqrs | 2 | **0.935141** | 2 | **0.979622** | 1 | 0.330808 | 1 | 0.446282 |
| matched_filter | 3 | **0.831949** | 4 | 0.670094 | 4 | 0.131714 | 4 | 0.078983 |
| two_average | 4 | 0.665419 | 3 | **0.820191** | 5 | 0.093727 | 3 | 0.140282 |
| christov | 5 | 0.658170 | 1 | **0.987552** | 2 | 0.218265 | 2 | 0.194806 |
| hamilton | 6 | 0.416173 | 6 | 0.260928 | 6 | 0.044460 | 7 | 0.015921 |
| swt | 7 | 0.189599 | 8 | 0.092911 | 8 | 0.033991 | 6 | 0.021340 |
| pan_tompkins | 8 | 0.176640 | 7 | 0.231381 | 7 | 0.044390 | 8 | 0.004288 |

**TABLE S35:**
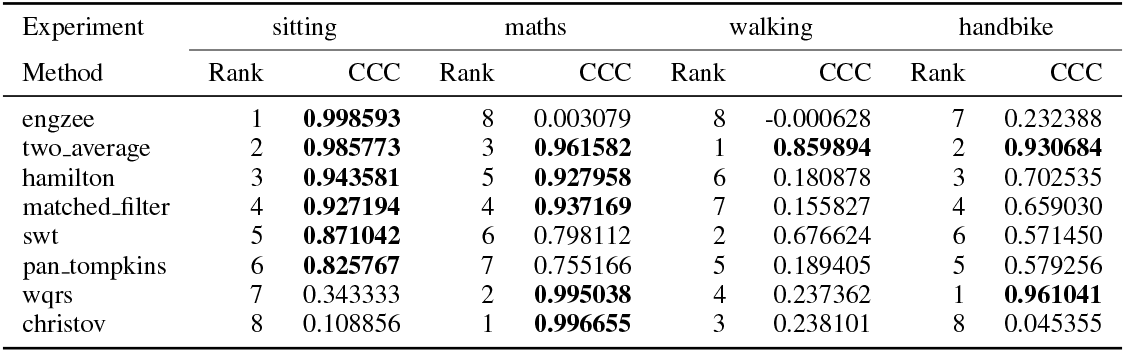
Ranking loose cables setup - Metric IQRNN.

| Experiment | sitting |  | maths |  | walking |  | handbike |  |
| --- | --- | --- | --- | --- | --- | --- | --- | --- |
| Method | Rank | CCC | Rank | CCC | Rank | CCC | Rank | CCC |
| engzee | 1 | **0.998593** | 8 | 0.003079 | 8 | -0.000628 | 7 | 0.232388 |
| two_average | 2 | **0.985773** | 3 | **0.961582** | 1 | **0.859894** | 2 | **0.930684** |
| hamilton | 3 | **0.943581** | 5 | **0.927958** | 6 | 0.180878 | 3 | 0.702535 |
| matched_filter | 4 | **0.927194** | 4 | **0.937169** | 7 | 0.155827 | 4 | 0.659030 |
| swt | 5 | **0.871042** | 6 | 0.798112 | 2 | 0.676624 | 6 | 0.571450 |
| pan_tompkins | 6 | **0.825767** | 7 | 0.755166 | 5 | 0.189405 | 5 | 0.579256 |
| wqrs | 7 | 0.343333 | 2 | **0.995038** | 4 | 0.237362 | 1 | **0.961041** |
| christov | 8 | 0.108856 | 1 | **0.996655** | 3 | 0.238101 | 8 | 0.045355 |

**TABLE S36:** Ranking loose cables setup - Metric LF.

| Experiment | sitting |  | maths |  | walking |  | handbike |  |
| --- | --- | --- | --- | --- | --- | --- | --- | --- |
| Method | Rank | CCC | Rank | CCC | Rank | CCC | Rank | CCC |
| pan_tompkins | 1 | **0.899065** | 2 | **0.956115** | 5 | 0.708830 | 8 | 0.125834 |
| two_average | 2 | **0.882706** | 5 | **0.89494** | 2 | **0.937494** | 1 | **0.900695** |
| swt | 3 | **0.857161** | 3 | **0.946041** | 1 | **0.971266** | 3 | 0.495127 |
| christov | 4 | **0.81944** | 1 | **0.959247** | 4 | 0.760662 | 2 | 0.759585 |
| hamilton | 5 | **0.805615** | 6 | **0.884232** | 3 | **0.936903** | 6 | 0.188718 |
| matched_filter | 6 | 0.696660 | 4 | **0.896769** | 7 | 0.489282 | 5 | 0.388615 |
| engzee | 7 | 0.588656 | 7 | **0.859338** | 8 | 0.488176 | 7 | 0.180855 |
| wqrs | 8 | 0.577444 | 8 | **0.814462** | 6 | 0.627607 | 4 | 0.431521 |

**TABLE S37:** Ranking loose cables setup - Metric HF.

| Experiment | sitting |  | maths |  | walking |  | handbike |  |
| --- | --- | --- | --- | --- | --- | --- | --- | --- |
| Method | Rank | CCC | Rank | CCC | Rank | CCC | Rank | CCC |
| hamilton | 1 | **0.802102** | 2 | 0.656655 | 3 | 0.322536 | 6 | 0.129962 |
| christov | 2 | 0.712909 | 3 | 0.648564 | 7 | 0.122749 | 1 | 0.678885 |
| matched_filter | 3 | 0.711363 | 7 | 0.180883 | 6 | 0.161254 | 3 | 0.305131 |
| swt | 4 | 0.575770 | 5 | 0.418717 | 2 | 0.337541 | 4 | 0.183217 |
| pan_tompkins | 5 | 0.570882 | 6 | 0.381738 | 4 | 0.247781 | 8 | 0.038249 |
| two_average | 6 | 0.414628 | 4 | 0.566373 | 1 | 0.370833 | 2 | 0.568125 |
| wqrs | 7 | 0.293930 | 1 | **0.862088** | 5 | 0.163661 | 7 | 0.066522 |
| engzee | 8 | 0.205098 | 8 | 0.131795 | 8 | -0.007703 | 5 | 0.143155 |

**TABLE S38:**
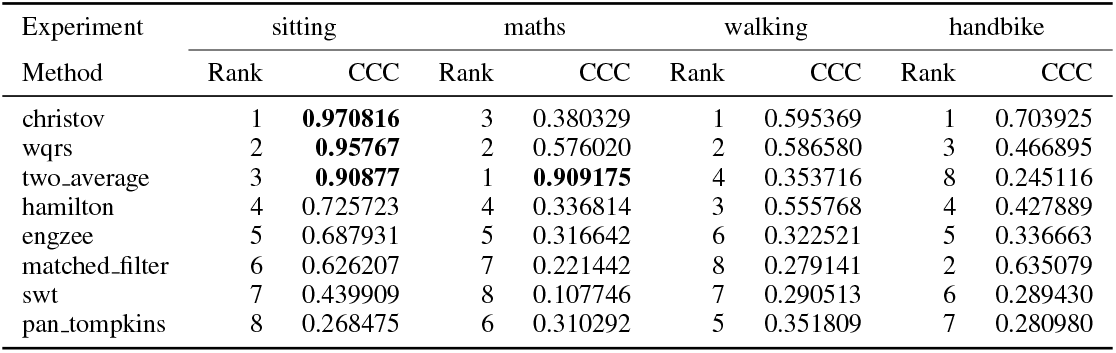
Ranking loose cables setup - Metric LFHF.

**TABLE S39:**
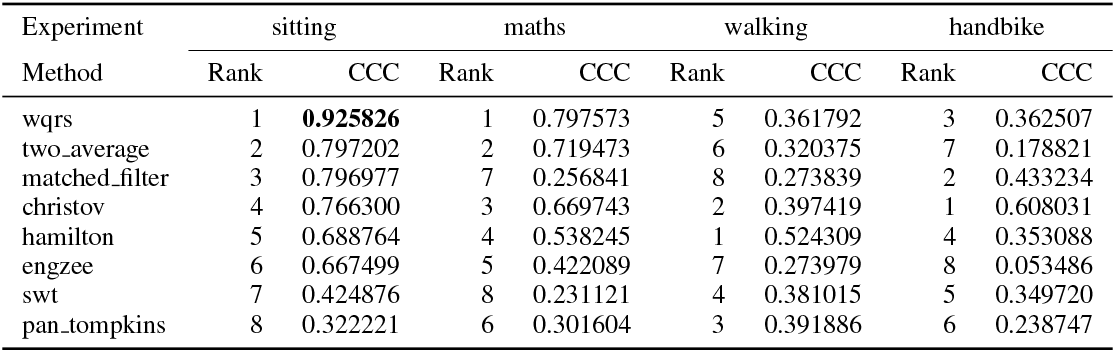
Ranking loose cables setup - Metric LFn.

| Experiment | sitting |  | maths |  | walking |  | handbike |  |
| --- | --- | --- | --- | --- | --- | --- | --- | --- |
| Method | Rank | CCC | Rank | CCC | Rank | CCC | Rank | CCC |
| wqrs | 1 | **0.925826** | 1 | 0.797573 | 5 | 0.361792 | 3 | 0.362507 |
| two_average | 2 | 0.797202 | 2 | 0.719473 | 6 | 0.320375 | 7 | 0.178821 |
| matched_filter | 3 | 0.796977 | 7 | 0.256841 | 8 | 0.273839 | 2 | 0.433234 |
| christov | 4 | 0.766300 | 3 | 0.669743 | 2 | 0.397419 | 1 | 0.608031 |
| hamilton | 5 | 0.688764 | 4 | 0.538245 | 1 | 0.524309 | 4 | 0.353088 |
| engzee | 6 | 0.667499 | 5 | 0.422089 | 7 | 0.273979 | 8 | 0.053486 |
| swt | 7 | 0.424876 | 8 | 0.231121 | 4 | 0.381015 | 5 | 0.349720 |
| pan_tompkins | 8 | 0.322221 | 6 | 0.301604 | 3 | 0.391886 | 6 | 0.238747 |

**TABLE S40:** Ranking loose cables setup - Metric HFn.

| Experiment | sitting |  | maths |  | walking |  | handbike |  |
| --- | --- | --- | --- | --- | --- | --- | --- | --- |
| Method | Rank | CCC | Rank | CCC | Rank | CCC | Rank | CCC |
| wqrs | 1 | **0.938161** | 1 | **0.878029** | 6 | 0.351714 | 4 | 0.278727 |
| hamilton | 2 | **0.913205** | 3 | **0.806907** | 2 | 0.524971 | 6 | 0.241423 |
| christov | 3 | **0.90781** | 4 | 0.736257 | 5 | 0.431526 | 1 | 0.540454 |
| two_average | 4 | **0.873838** | 2 | **0.809859** | 3 | 0.503703 | 7 | 0.185865 |
| matched_filter | 5 | **0.854991** | 8 | 0.415271 | 8 | 0.282450 | 2 | 0.412928 |
| engzee | 6 | 0.795775 | 6 | 0.533268 | 7 | 0.313248 | 8 | 0.060043 |
| swt | 7 | 0.783202 | 7 | 0.470260 | 4 | 0.499574 | 3 | 0.360251 |
| pan_tompkins | 8 | 0.680921 | 5 | 0.635104 | 1 | 0.574723 | 5 | 0.269342 |

**TABLE S41:** Ranking loose cables setup - Metric LnHF.

| Experiment | sitting |  | maths |  | walking |  | handbike |  |
| --- | --- | --- | --- | --- | --- | --- | --- | --- |
| Method | Rank | CCC | Rank | CCC | Rank | CCC | Rank | CCC |
| christov | 1 | **0.940616** | 4 | **0.816478** | 5 | 0.395774 | 1 | 0.774384 |
| two_average | 2 | 0.799493 | 1 | **0.885184** | 1 | 0.589140 | 4 | 0.360819 |
| hamilton | 3 | 0.791002 | 2 | **0.82921** | 2 | 0.582415 | 5 | 0.312152 |
| matched_filter | 4 | 0.763808 | 8 | 0.508666 | 7 | 0.263051 | 3 | 0.408452 |
| wqrs | 5 | 0.743609 | 3 | **0.824249** | 3 | 0.465389 | 6 | 0.284667 |
| swt | 6 | 0.638118 | 7 | 0.608933 | 6 | 0.363096 | 2 | 0.475706 |
| engzee | 7 | 0.607269 | 6 | 0.609647 | 8 | 0.088834 | 8 | 0.095081 |
| pan_tompkins | 8 | 0.466908 | 5 | 0.663671 | 4 | 0.414253 | 7 | 0.150432 |

**TABLE S42:**
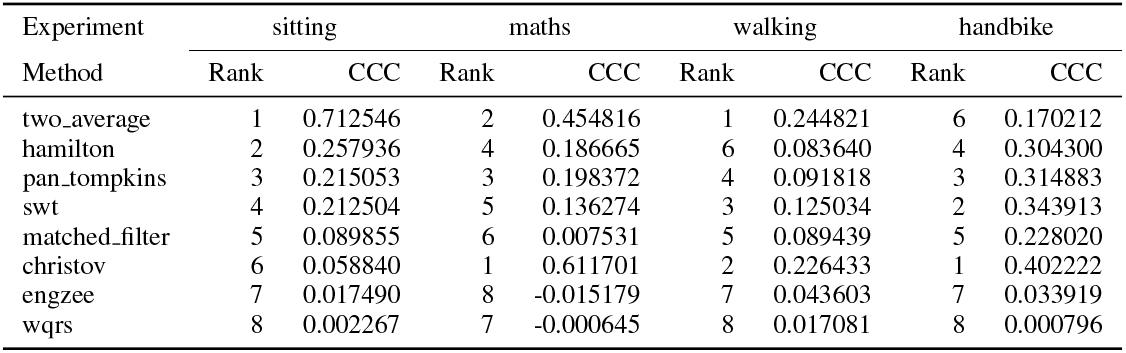
Ranking loose cables setup - Metric SD1.

**TABLE S43:**
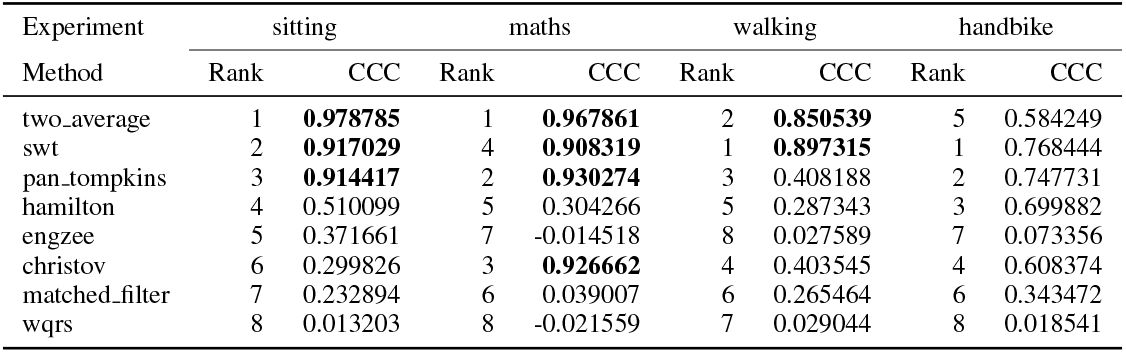
Ranking loose cables setup - Metric SD2.

| Experiment | sitting |  | maths |  | walking |  | handbike |  |
| --- | --- | --- | --- | --- | --- | --- | --- | --- |
| Method | Rank | CCC | Rank | CCC | Rank | CCC | Rank | CCC |
| two_average | 1 | **0.978785** | 1 | **0.967861** | 2 | **0.850539** | 5 | 0.584249 |
| swt | 2 | **0.917029** | 4 | **0.908319** | 1 | **0.897315** | 1 | 0.768444 |
| pan_tompkins | 3 | **0.914417** | 2 | **0.930274** | 3 | 0.408188 | 2 | 0.747731 |
| hamilton | 4 | 0.510099 | 5 | 0.304266 | 5 | 0.287343 | 3 | 0.699882 |
| engzee | 5 | 0.371661 | 7 | -0.014518 | 8 | 0.027589 | 7 | 0.073356 |
| christov | 6 | 0.299826 | 3 | **0.926662** | 4 | 0.403545 | 4 | 0.608374 |
| matched_filter | 7 | 0.232894 | 6 | 0.039007 | 6 | 0.265464 | 6 | 0.343472 |
| wqrs | 8 | 0.013203 | 8 | -0.021559 | 7 | 0.029044 | 8 | 0.018541 |

**TABLE S44:** Ranking loose cables setup - Metric SD1SD2.

| Experiment | sitting |  | maths |  | walking |  | handbike |  |
| --- | --- | --- | --- | --- | --- | --- | --- | --- |
| Method | Rank | CCC | Rank | CCC | Rank | CCC | Rank | CCC |
| two_average | 1 | 0.444132 | 2 | 0.488049 | 2 | 0.295028 | 6 | 0.063566 |
| christov | 2 | 0.311372 | 1 | 0.605134 | 1 | 0.437578 | 1 | 0.306289 |
| wqrs | 3 | 0.073234 | 6 | 0.093558 | 6 | 0.114542 | 7 | 0.054540 |
| engzee | 4 | 0.037893 | 3 | 0.225065 | 3 | 0.178233 | 8 | 0.012990 |
| matched_filter | 5 | 0.028861 | 5 | 0.096474 | 4 | 0.155106 | 2 | 0.154757 |
| swt | 6 | -0.032405 | 7 | 0.082441 | 8 | 0.076346 | 3 | 0.143349 |
| pan_tompkins | 7 | -0.049325 | 4 | 0.098250 | 7 | 0.110190 | 5 | 0.111505 |
| hamilton | 8 | -0.163027 | 8 | -0.000490 | 5 | 0.154065 | 4 | 0.143264 |

**TABLE S45:** Ranking loose cables setup - Metric SampEn.

| Experiment | sitting |  | maths |  | walking |  | handbike |  |
| --- | --- | --- | --- | --- | --- | --- | --- | --- |
| Method | Rank | CCC | Rank | CCC | Rank | CCC | Rank | CCC |
| two_average | 1 | **0.807298** | 2 | **0.814312** | 1 | 0.719380 | 1 | **0.802751** |
| christov | 2 | 0.747562 | 1 | **0.845186** | 2 | 0.359500 | 2 | 0.611742 |
| engzee | 3 | 0.602449 | 5 | 0.513601 | 3 | 0.264014 | 4 | 0.356524 |
| pan_tompkins | 4 | 0.398491 | 3 | 0.693708 | 8 | 0.034709 | 5 | 0.223837 |
| hamilton | 5 | 0.264666 | 7 | 0.152607 | 7 | 0.044104 | 3 | 0.468628 |
| wqrs | 6 | 0.199285 | 6 | 0.246586 | 6 | 0.099472 | 8 | 0.093417 |
| swt | 7 | 0.192423 | 8 | 0.057307 | 4 | 0.196044 | 6 | 0.184833 |
| matched_filter | 8 | 0.155287 | 4 | 0.560902 | 5 | 0.126716 | 7 | 0.158137 |

**TABLE S46:**
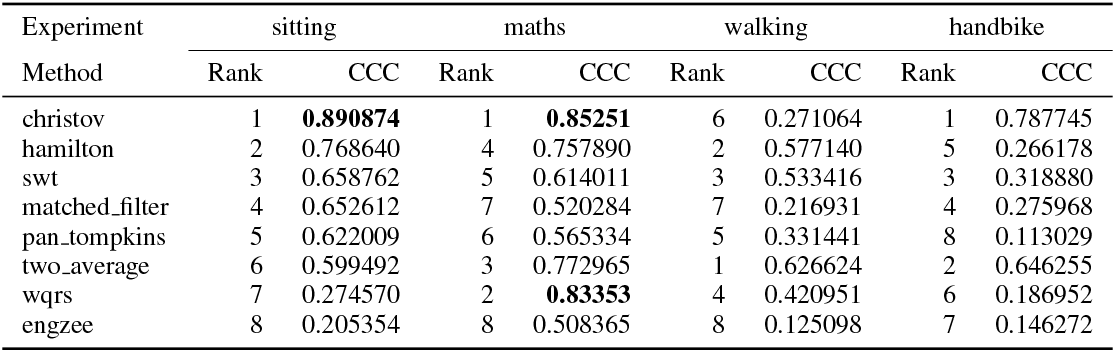
Ranking loose cables setup - Metric TP.

## Footnotes

1 An ectopic beat is a beat with an origin different than the sinoatrial node. The presence of these beats in the interval tachogram is considered an artifact and must be removed before computing an HRV metric [13], [15].

## Notes

This work was supported in part by Agencia Nacional de Investigación y Desarrollo (ANID): Grants BASAL AFB240002 (A.W.), BASAL FB210008 (M.O.), Anillo ACT210053, FONDECYT INICIACION 11241484 (M.O.), FONDECYT EXPLORACION 13240042 (M.O.), FONDECYT REGULAR 1231132 (A.W.).

### Competing Interest Statement

The authors have declared no competing interest.

